# Characterising antibody kinetics from multiple influenza infection and vaccination events in ferrets

**DOI:** 10.1101/411751

**Authors:** James A. Hay, Karen Laurie, Michael White, Steven Riley

**Affiliations:** MRC Centre for Global Infectious Disease Analysis, Department of Infectious Disease Epidemiology, Imperial College, London, UK; WHO Collaborating Centre for Reference and Research on Influenza, Peter Doherty Institute for Infection and Immunity, Melbourne, Australia; Seqirus, 63 Poplar Road, Parkville Victoria 3052, Australia; Malaria: Parasites and Hosts, Department of Parasites and Insect Vectors, Institut Pasteur, Paris, France

## Abstract

The strength and breadth of an individual’s antibody repertoire are important predictors of their response to influenza infection or vaccination. Although progress has been made in understanding qualitatively how repeated exposures shape the antibody mediated immune response, quantitative understanding remains limited. We developed a set of mathematical models describing short-term antibody kinetics following influenza infection or vaccination and fit them to haemagglutination inhibition (HI) titres from 5 groups of ferrets which were exposed to different combinations of trivalent inactivated influenza vaccine (TIV with or without adjuvant), A/H3N2 priming inoculation and post-vaccination A/H1N1 inoculation. We fit models with various immunological mechanisms that have been empirically observed but are yet to be included in mathematical models of antibody landscapes, including titre ceiling effects, antigenic seniority and exposure-type specific cross reactivity. Based on the parameter estimates of the best supported models, we describe a number of key immunological features. We found quantifiable differences in the degree of homologous and cross-reactive antibody boosting elicited by different exposure types. Infection and adjuvanted vaccination generally resulted in strong, broadly reactive responses whereas unadjuvanted vaccination resulted in a weak, narrow response. We found that the order of exposure mattered: priming with A/H3N2 improved subsequent vaccine response, and the second dose of adjuvanted vaccination resulted in substantially greater antibody boosting than the first. Either antigenic seniority or a titre ceiling effect were included in the two best fitting models, suggesting that a mechanism describing diminishing antibody boosting with repeated exposures improved the predictive power of the model. Although there was considerable uncertainty in our estimates of antibody waning parameters, our results suggest that both short and long term waning were present and would be identifiable with a larger set of experiments. These results highlight the potential use of repeat exposure animal models in revealing short-term, strain-specific immune dynamics of influenza.

**Author summary:** Despite most individuals having some preexisting immunity from past influenza infections and vaccinations, a significant proportion of the human population is infected with influenza each year. Predicting how an individual’s antibody profile will change following exposure is therefore useful for evaluating which populations are at greatest risk and how effective vaccination strategies might be. However, interpretation of antibody data from humans is complicated by immunological interactions between all previous, unobserved exposures in an individual’s life. We developed a mathematical model to describe short-term antibody kinetics that are important in building an individual’s immune profile but are difficult to observe in human populations. We validated this model using antibody data from ferrets with known, varied infection and vaccination histories. We were able to quantify the independent contributions of various exposures and immunological mechanisms in generating observed antibody titres. These results suggest that data from experimental systems may be included in models of human antibody dynamics, which may improve predictions of vaccination strategy effectiveness and how population susceptibility changes over time.

## Introduction

Natural infection with influenza stimulates a complex and multifaceted immune response to neutralise and clear the infection. [1] The adaptive immune response is of particular interest for seasonal epidemic and pandemic preparedness, as responses from previous exposures provide some long-term protection against reinfection and disease via antibody and T-cell mediated immunity. [2,3] Focusing on the adaptive immune response is also advantageous because it can (i) be induced in advance of an epidemic through vaccination and (ii) be measured and compared against correlates of protection to improve public health forecasting. [4–7] However, influenza is an antigenically variable virus and undergoes continual antigenic drift, whereby mutations in immunodominant epitopes are selected by immunological pressure, allowing influenza lineages to escape population herd immunity. [8–10] This results in the continual waning of long-term immunity as antibodies effective against past strains fail to neutralise novel variants. [11] The current strategy for combating antigenic drift is to regularly update the seasonal influenza vaccine to better represent circulating strains, resulting in an arms race between virus and vaccine formulation. This approach is considered sub-optimal given the failure rate of matching and manufacturing the vaccine strain in advance of the epidemic period. [12,13] Consequently, there has been a recent push towards a universal influenza vaccination strategy, either through new vaccines or improved strength and breadth of immunity using existing technologies. [14,15]

Whilst there is some cross-reactivity and cross-protection within influenza A virus subtypes and within influenza B virus lineages, humans experience numerous infections over their lives. [16–18] Each successive influenza exposure, which may be vaccination or infection, can strengthen the available repertoire of T and B cells which target epitopes on circulating and previously encountered strains. [19,20] In the humoral response, this occurs by boosting antibodies produced by preexisting long-lived plasma cells and activated memory B cells (MBCs), and through generating a novel B cell response targeting unrecognised epitopes. [21–23] Given that individuals experience repeated infections and vaccinations from antigenically varied influenza viruses, interpreting the composition of an observed antibody response is confounded by the complex interaction of an individual’s preexisting immunity, or immune history, with the infecting virus. [24–27]

A large body of experimental and observational work exists describing the contribution of immune histories to observed influenza susceptibility profiles, antibody landscapes and vaccination responses, often under the terms “original antigenic sin” or “antigenic seniority”. [1,28–33] Furthermore, next generation assays to characterise antibody diversity and B-cell identity have provided a detailed understanding of immune dynamics, including short term immune kinetics, duration of the humoral response, and immunodominance of different antigenic sites. [34–38] However, few studies have integrated these mechanisms into quantitative frameworks which can be used to explain and predict serological data from human populations, which often rely on simpler and less finely resolved assays. [18,22,39,40] Human models for influenza are also difficult to interpret due to highly varied and unobserved exposure histories. Animal models, in particular ferrets, have therefore been used to generate much of our understanding of influenza immunology due to opportunity for intensive observation and control. [34,41–46] Here, we exploit the experimental flexibility and transparency of a ferret model to find evidence for and quantify multiple immunological mechanisms that may be important in characterizing antibody landscapes generated from complex exposure histories, yet are observable using only routine antibody assays. Quantifying short term mechanisms in a ferret system might reveal patterns that could be used to improve the predictability and interpretation of human antibody landscapes following exposure. [47]

We developed a mathematical model of antibody boosting and biphasic waning to describe antibody kinetics using previously published antibody titre data from a group of ferrets with varied but known exposure histories. [41] Previous models of antibody kinetics have often focused on the response to a single immunogen following one exposure. [48–50] Here, we take into account previously described immunological phenomena to describe cross-reactive antibody titres arising from varied exposure histories. These phenomena included exposure type-specific homologous and cross-reactive antibody kinetics, the role of priming on subsequent vaccination, titre-dependent antibody boosting (or titre ceiling effects) and reduced antibody boosting with each subsequent exposure (antigenic seniority). [31] By fitting models with various combinations of these mechanisms to haemagglutination inhibition (HI) titre data from ferrets, we sought to identify immunological mechanisms that are important in describing observed antibody profiles arising from multiple exposures. Parameter estimates from these model fits allowed us to quantify the impact of prior infection and adjuvant inclusion on antibody levels following vaccination and to compare homologous and cross reactive boosting profiles of different exposure types. [51–55]

## Materials & Methods

### Study Data

Antibody titre data were obtained from a previously published ferret study. [41] The experimental protocol was originally designed to reflect different possible human infection and vaccination histories at the time of the 2009 pandemic. Here we present a secondary analysis of these data, with the intention of characterizing underlying immunological processes. Although these experiments were not designed specifically to power our mathematical model, they provide an example of how such studies may be designed to parameterise immunological parameters of interest.

Briefly, five experimental groups each of three ferrets underwent different combinations of infection with seasonal influenza A and/or vaccination with Northern and Southern Hemisphere trivalent inactivated influenza vaccine (TIV), with or without Freund’s incomplete adjuvant, over the course of 70 days (Table 1). Serum samples were collected at days 0, 21, 37, 49 and 70 from all ferrets (Fig 1). HI titres were used to determine antibody titres to each infection and TIV strain. Dilution plates with 12 wells were used, such that the highest possible recorded dilution was 1:40960, and the lowest detectable titre was 1:20. Undetectable titres were recorded as <1:20. All analyses here were carried out using log titres, defined as *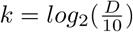*, where *D* was the recorded dilution. Observed log titres were therefore assigned values between 0 and 12, where < 1 : 20 = 0, 1 : 20 = 1 and ≥ 1 : 40960 = 12.

**Table 1.** Description of experimental protocol.

| Group | Infection with A/Panama/2007/99 (H3N2) | Immunisation with S.H TIV 2008* | Immunisation with N.H TIV 2007/2008** | Infection with A/Fukushima/141/06 (H1N1) | Number of ferrets |
| --- | --- | --- | --- | --- | --- |
| Group A | No | Yes (no adjuvant) | Yes (no adjuvant) | Yes | 3 |
| Group B | No | Yes (with adjuvant) | Yes (with adjuvant) | Yes | 3 |
| Group C | Yes | Yes (no adjuvant) | Yes (no adjuvant) | Yes | 3 |
| Group D | Yes | Yes (with adjuvant) | Yes (with adjuvant) | Yes | 3 |
| Group E | Yes | No | No | Yes | 3 |

\*Southern Hemisphere (S.H.) TIV 2008: A/Solomon Islands/3/2006 (H1N1), A/Brisbane/10/2007 (H3N2), B/Brisbane/3/2007
\*\*Northern Hemisphere (N.H.) TIV 2007/2008: A/Solomon Islands/3/2006 (H1N1), A/Wisconsin/67/2005 (H3N2), B/Malaysia/2506/2004

**Fig 1.**
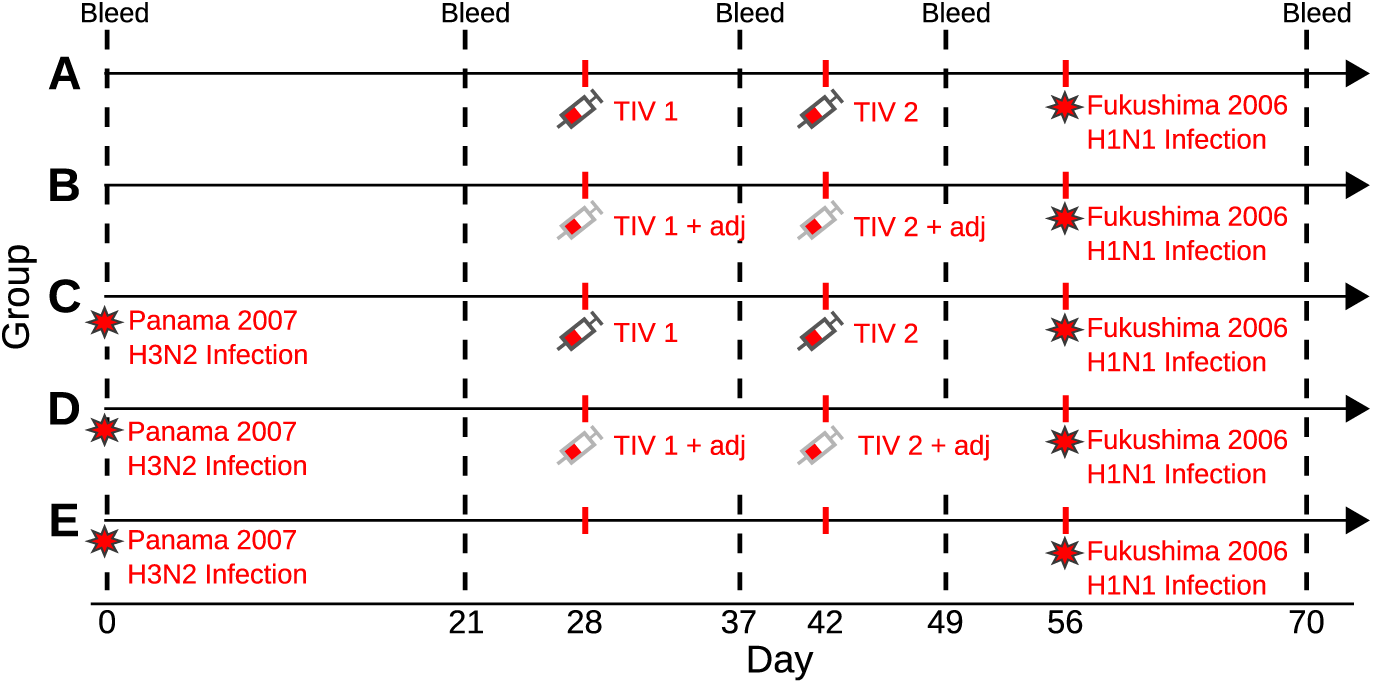
Summary of experimental protocol. Days since first event are shown on the x-axis, with the 5 groups shown as rows. Red stars represent infection with either A/Panama/2007/1999 (H3N2) or A/Fukushima/141/2006 (H1N1). Red syringes represent vaccination with either Southern Hemisphere TIV 2008 (TIV 1) or Northern Hemisphere TIV 2007/2008 (TIV 2) with (grey border) or without (black border) adjuvant. Vertical, dashed black lines represent times of blood sample collection, providing HI titres against each of the vaccination and infection strains at that time point.

Full adult doses of human TIV were used in groups A, B, C and D. The first vaccination (Southern Hemisphere 2008 TIV) contained A/Solomon Islands/3/2006 (H1N1), A/Brisbane/10/2007 (H3N2) and B/Brisbane/3/2007, administered at day 28 (TIV 1). The second vaccination (Northern Hemisphere 2007/2008 TIV) contained A/Solomon Islands/3/2006 (H1N1), A/Wisconsin/67/2005 (H3N2) and B/Malaysia/2506/2004, administered at day 42 (TIV 2). Vaccines used in groups B and D were emulsified in an equal volume of Freund’s incomplete adjuvant immediately before administration (TIV 1/2 + adjuvant). All vaccines contained 15*µ*g of HA of each strain, and were delivered to sedated animals intramuscularly in the quadriceps muscles of both hind legs. Infections were carried out by dropwise intranasal challenges with 10^3.5^ 50% tissue culture infectious doses (TCID_50_) in 0.5 mL with A/Panama/2007/1999 (H3N2) in groups C, D and E, and with A/Fukushima/141/2006 (H1N1) in all groups.

### Models of antibody kinetics

The mathematical model describes the kinetics of homologous and heterologous antibody titres following exposure. Fig 2 depicts the example of an individual becoming infected and later vaccinated, though the model may characterise any sequence of exposures. Conceptually similar mathematical models of boosting followed by biphasic waning have been used previously to describe antibody secreting cell (ASC) and antibody kinetics. [48,50,56]

**Fig 2.**
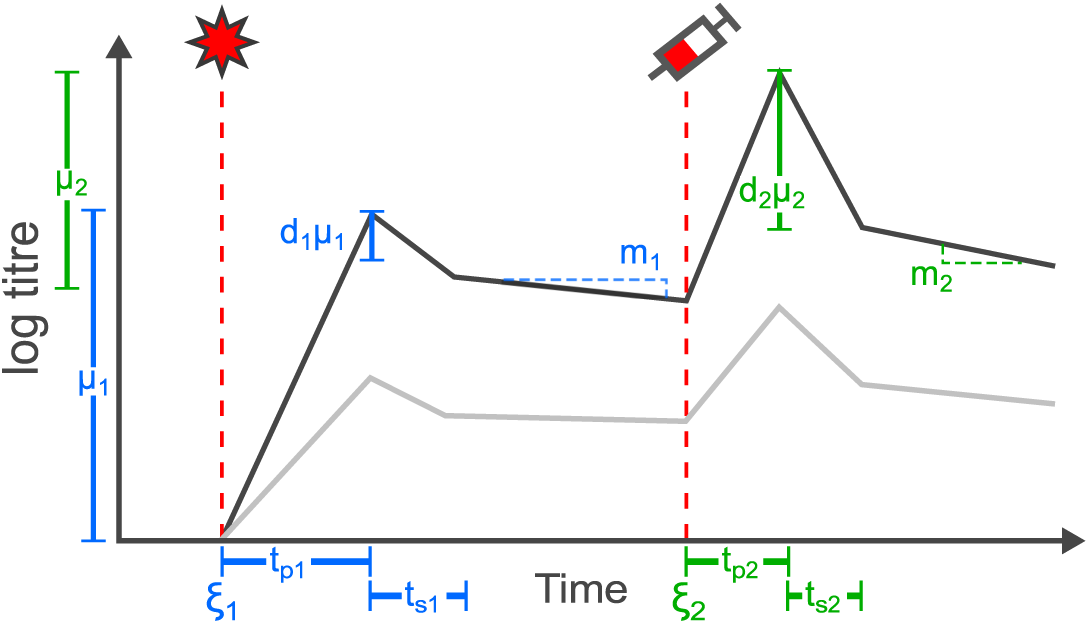
Base model. Schematic showing the relationship between model parameters and antibody kinetics over time. Black line shows antibody titres effective against one immunogen. Grey line highlights how antibody titres to a different influenza immunogen (that is less antigenically similar to the exposure immunogens than the black line) develop in parallel driven by cross-reactive antibodies. After each exposure, antibody levels undergo linear boosting on a log scale followed by an initial, short waning phase and then a slower, long-term waning phase. This example demonstrates two exposures, initially with infection (star symbol) and subsequently with vaccine (syringe symbol), where antibody dynamics are governed by a set of parameters depending on the exposure type. Note that the y-axis is on a log scale and all observations are discrete and taken as the floor value. Model parameters are described in S1 Table.

After an infection (start at time *ξ*_1_), homologous antibody titres undergo boosting rising linearly (on the log scale) to a peak after time *t*_*p*1_, ignoring any delay between exposure and the start of antibody production. Titres then quickly drop by a fixed proportion, *d*_1_, over time *t*_*s*1_, representing the initial short-term waning phase as free antibodies and early short-lived ASCs begin to decay following clearance of the initial antigen dose. [32] Antibody waning then switches to a constant rate *m*_1_ (log titre units lost per day) for the remainder of time (representing the population of persistent ASCs) until subsequent vaccination (syringe at time *ξ*_2_), when antibody dynamics become dominated by a new set of boosting and waning parameters. We did not include a third, steady state phase due to the short time frame of these experiments. [57] Antibodies effective against heterologous strains experience boosting and biphasic waning in proportion to the exposure strain, with the proportion dependent on the antigenic distance between the measured and exposure strains.

We then built on this base model to incorporate additional immunological mechanisms that are important in describing antibody boosting and waning. These included: biphasic or monophasic antibody waning; exposure-type specific or type non-specific cross-reactivity; antigenic seniority; the impact of priming infection on subsequent vaccine response; and titre-dependent boosting. We considered models with different numbers of exposure types: either 3 (infection, TIV, TIV + adjuvant) or 6 (priming infection, secondary infection, initial TIV, secondary TIV, initial TIV + adjuvant, secondary TIV + adjuvant). The base boosting and waning model remains the same across model variants, but these mechanisms add complexity to the boosting parameter, *µ*, and link different exposures with common parameters. A full description of each of these mechanisms and their implementation is described in S1 Supporting Protocol.

We fit each of the 64 potential model variants in a Bayesian framework using parallel-tempering Markov chain Monte Carlo (PT-MCMC) to estimate the posterior medians and 95% credible intervals (CI) of all free model parameters. For each model, we ran 3 chains each for 5000000 iterations. Where the effective sample size (ESS) was < 200 or the Gelman-Rubin diagnostic 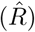 was < 1.1 for any estimated parameter (calculated using the ‘coda’ R package), we ran 5 chains each for 10000000 iterations and obtained upper 95% confidence intervals for 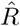 of < 1.1 for all estimated parameters presented here. [58] ESS and 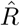 estimates for all parameters are provided in S5 Table.

We then performed a model comparison analysis using Pareto-smoothed importance sampling leave-one-out cross-validation (PSIS-LOO) with the ‘loo’ R package. [59,60] Briefly, the purpose of this analysis was to compare the expected log pointwise predictive density (ELPD) of different model fits to compare their out-of-sample prediction accuracy. Comparing ELPD estimates serves a similar purpose to comparing other information criteria, where a lower ELPD suggests greater predictive power penalised by model complexity. Results shown in the main text are from the most complex model (most free parameters) variant with *δ*ELPD<1 compared to the lowest ELPD. Parameter estimates from all model variants with a *δ*ELPD<20 are shown in Figures S5-12. Posterior parameter estimates are shown as medians and 95% credible intervals (CIs). Further details of the model fitting and comparison are described in S1 Supporting Protocol. All code and data are available as an R package at https://github.com/jameshay218/antibodyKinetics.

## Results

### Antibody kinetics following a single exposure support biphasic waning

To validate our boosting and biphasic waning model for a single exposure, we fit the base model to HI titres against A/Panama/2007/1999 (H3N2) from group E alone (Fig 3). These data in isolation reflect a typical antibody trajectory following exposure to a single immunogen and measurement of antibodies against it. [33,48] The models with biphasic waning (both with estimated long term waning rate, *m*, and fixed *m* = 0) were better supported than the models with monophasic waning or no waning (ELPD-20.6 (standard error (SE), 3.35) and −20.5 (SE, 3.78) compared to −23.4 (SE, 3.71) and −28.1 (SE, 3.48) respectively), although we note that these differences are small with respect to the standard error of the ELPD estimates. The biphasic waning models with estimated long-term waning *m* and fixed long-term waning *m* = 0 had a difference in ELPD of <1, suggesting that both models had similar predictive performance. Overall, these results suggest that the model with monophasic waning is justified over the version with no waning, and that the biphasic waning model is better still than the monophasic waning model. Posterior estimates for model parameters were: *µ* = 9.91 (median, 95% CI 7.08-12.7); *d* = 0.551 (median, 95% CI 0.183-0.695); *t*_*s*_ = 19.5 days (median, 95% CI 6.39-27.4 days); *m* = 0.0414 (median, 95% CI 0.00405-0.103).

**Fig 3.**
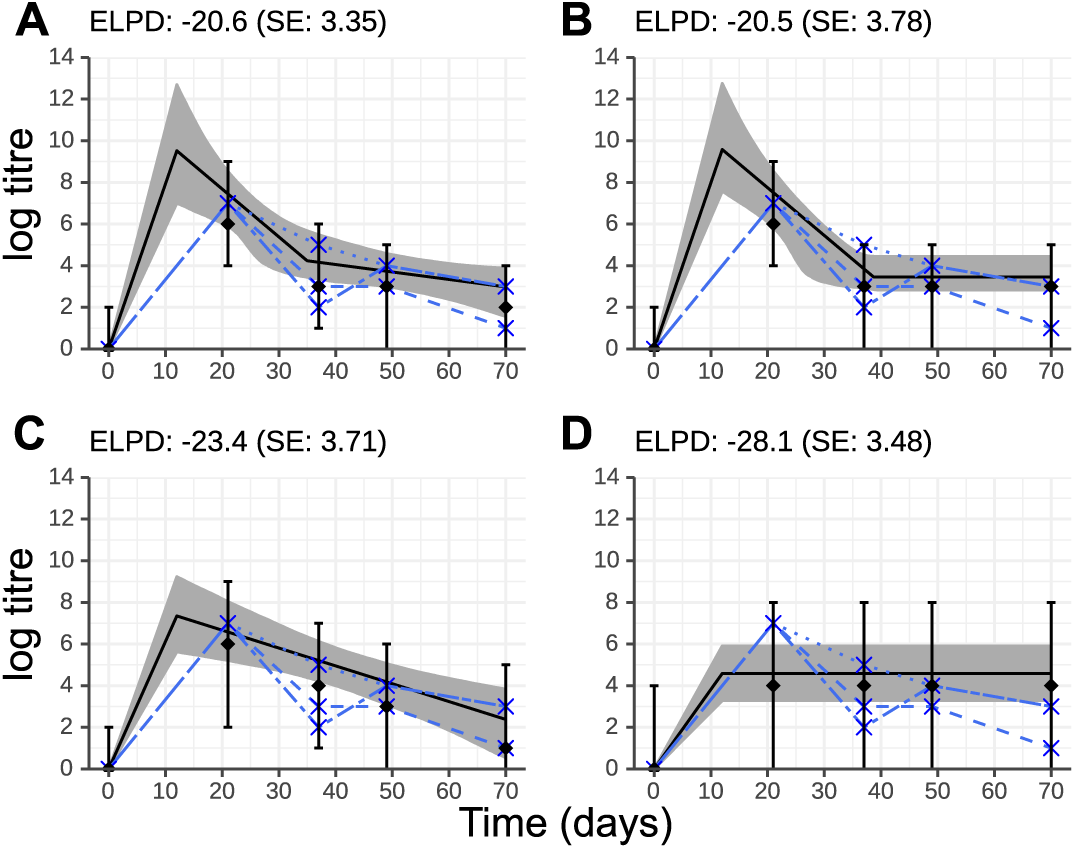
Comparison of four model fits to data from three ferrets following exposure to a single immunogen. All ferrets were infected with A/Panama/2007/99 (H3N2) at day 0. Y-axis shows log HI titre against A/Panama/2007/99 (H3N2). Solid black line and grey region show best-fit model trajectory and 95% credible intervals (CI) of latent antibody titre. Black diamond and error bars show median and 95% CI of model predicted observations. Blue crosses and dashed lines show observed log HI titre for the three individual ferrets. Figure titles show estimated expected log-predictive density (ELPD) and corresponding standard error (SE). A: Biphasic waning; B: biphasic waning with a fixed long-term waning rate of *m* = 0; C: monophasic waning with *t*_*s*_ = *d* = 0; D; no short or long term waning with *m* = *t*_*s*_ = *d* = 0.

### Variation in antibody kinetics driven by different exposure histories

Overall, ferrets that received more frequent and immunogenic exposures achieved the highest, most broadly reactive and long-lived antibody titres. The full data show substantial variation in observed antibody titres across the groups driven by different exposure types and combinations. Following two doses of unadjuvanted TIV, ferrets achieved only modest increases in titres against the vaccine strains (Fig 4A), with 2 out of 3 ferrets failing to generate H3N2 titres that persisted past day 37. The addition of an adjuvant resulted in increased and persistent titres against the vaccine strains in all ferrets by day 49. Titres against A/Fukushima/141/2006 (H1N1), which is antigenically similar to A/Solomon Islands/3/2006 (H1N1), were also increased at this timepoint (Fig 4B). Similarly, priming infection resulted in higher and long-lived titres to the vaccine strains and A/Fukushima/141/2006 (H1N1) relative to ferrets in the unprimed, unadjuvanted TIV protocol (Fig 4C). Observed titres at day 21 against A/Panama/2007/1999 (H3N2) were consistently high following priming infection in groups C-E, with one ferret in each of groups C and E also experiencing some boosting of antibodies against the other H3N2 strains. All ferrets were infected with A/Fukushima/141/2006 (H1N1) at day 56, leading to elevated titres to both H1N1 strains by day 70 in all ferrets.

**Fig 4.**
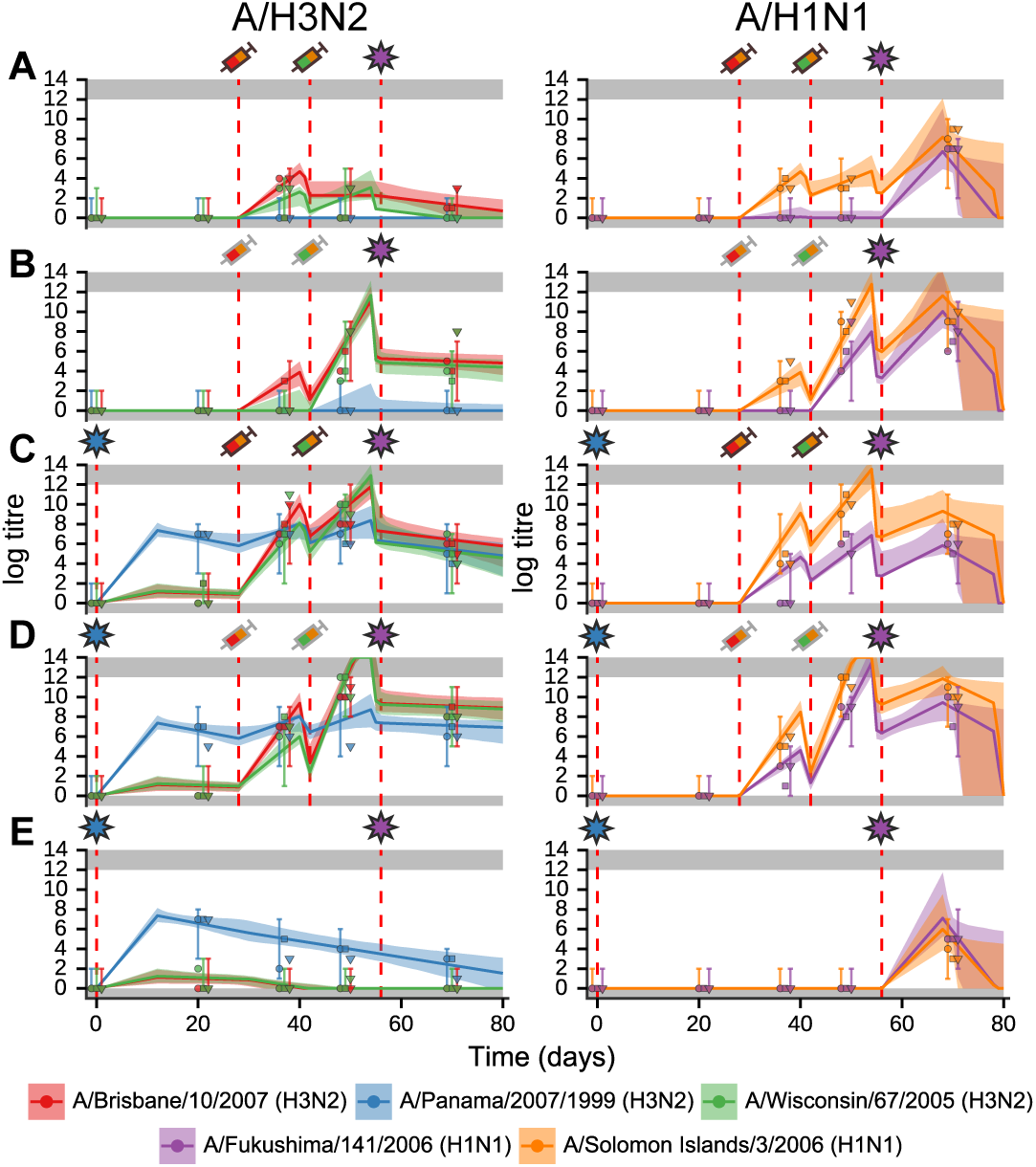
Model trajectory fits. coloured lines represent the best-fit trajectories of antibody kinetics following exposure. Coloured shaded regions show 95% credible intervals of the model fit. coloured points show observed discrete log antibody titres by HI assay for each of the three individual ferrets in each group. Gray shaded regions show the upper and lower limits of detection in the assay. For the same ferrets, titres to A/H3N2 strains are shown in the left column and A/H1N1 strains in the right column. Red dashed lines show times of exposures. Groups A-E correspond to descriptions in Fig 1. Exposure events are as described in Table 1. Symbols above each subplot: star represents infection; syringe represents TIV; and syringe with gray border represents TIV + adjuvant. Symbols are coloured based on their formulation. TIV 1 contained the following influenza A strains: A/Solomon Islands/3/2006 (H1N1) and A/Brisbane/10/2007 (H3N2). TIV 2 contained: A/Solomon Islands/3/2006 (H1N1) and A/Wisconsin/67/2005 (H3N2).

### Model comparison results

The top two model variants had ELPD estimates of −412.1 (SE, 21.4) and −412.6 (SE, 20.7) respectively. Both of these models included: a role for priming infection in increasing subsequent vaccine response; different boosting profiles between vaccination and infection; different boosting profiles with adjuvant versus without adjuvant; and biphasic antibody waning. The model with lowest the ELPD had 30 free parameters and also included titre-dependent boosting, no antigenic seniority, and no exposure type-specific cross reactivity. The other model had 33 free parameters and did not include titre-dependent boosting, but did include antigenic seniority and exposure type-specific cross reactivity. Fig 4 shows the latter (more complex) model variant fitted to the data. Parameter estimates for these two models are shown in S3 Table. The remainder of the results refer to the latter model with more free parameters.

Overall, ELPD estimates ranged from −412.1 (SE, 21.4) in the highest ranked model to −543.6 (SE, 21.8) in the lowest ranked model. The most simple model with 8 free parameters was the third lowest ranked model (ELPD −539.0 (SE, 21.9)), whereas the most complex model with 35 free parameters was the the 7th highest ranked model (ELPD −417.0 (SE, 21.2)). We note that some of the simpler model variants may have similar predictive performance to the best fitting, complex model and may therefore be more suitable in a general predictive application. However, our aim was not to predict unseen data here but rather to quantify immunological mechanisms.

As a crude measure of mechanism importance, we performed Pseudo-Bayesian model averaging (Pseudo-BMA+) to estimate the relative weights of each model variant and thereby weights of models with a particular mechanism relative to models without. [61] Although comparison of variable importance using information criteria must be interpreted with caution (for example, changing the sample size or experimental protocol may change the results), Pseudo-BMA+ serves as a rough estimate of which mechanisms are most important in explaining these data. [62] Variable weights were: 1.00 for the presence of priming; 0.999 for the presence of 6 exposure types; 0.836 for the presence of biphasic waning; 0.579 for the presence of titre dependent boosting; 0.572 for the presence of type specific cross reactivity; and 0.406 for the presence of antigenic seniority. The top two models had Pseudo-BMA+ weights of 0.331 and 0.303, with a drop off to the third model with a weight of 0.0977. The top two models included only titre-dependent boosting and antigenic seniority respectively, suggesting that inclusion of at least one of these mechanisms improved predictive performance. The consistency of parameter estimates across the best fitting model variants is demonstrated in Figures S5-12.

### Comparison of homologous boosting by exposure type

The level of homologous boosting resulting from priming infection (Infection 1) and secondary infection (Infection 2) appeared to be similar. Similar estimates were obtained for *µ* from both infections (Fig 5i), suggesting that the antibody response following priming infection appeared to be persistent. We inferred that antibody titres fell only marginally following the initial waning phase (*µ*(1 – *d*), Fig 5ii). The antibody waning rate was not identifiable for secondary infection due to the lack of observations following this exposure. We found evidence for only low levels of homologous antibody boosting following both initial and secondary doses of unadjuvanted TIV (TIV 1 and TIV 2) that quickly waned to near undetectable levels during the initial waning phase.

**Fig 5.**
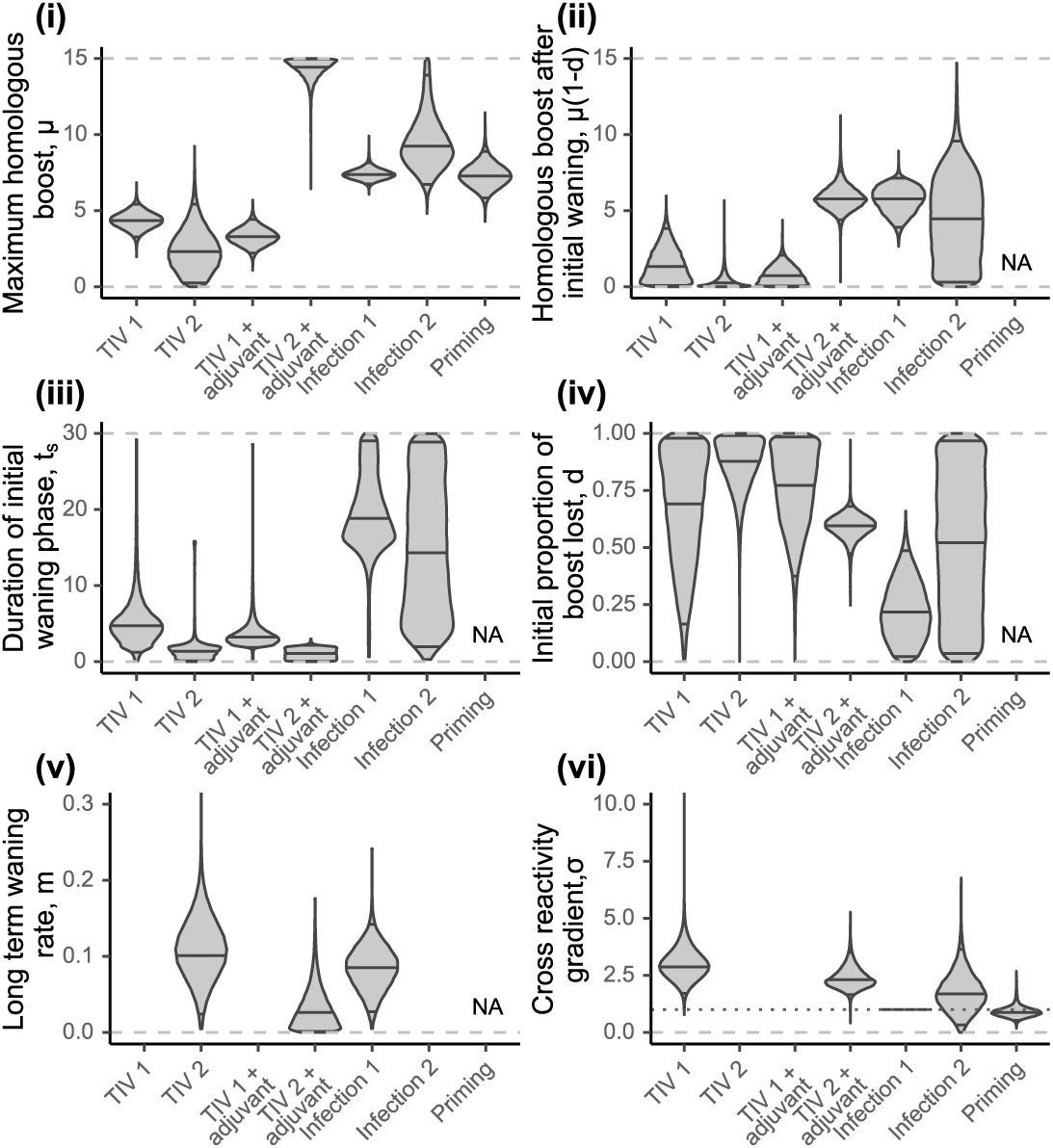
Estimated model parameters. Violin plots showing estimated posterior densities with medians and 95% credible intervals marked as horizontal black lines. Violin plots are similar to boxplots, but show the full probability density with some smoothing through a kernel density estimator. Dashed gray lines show bounds on uniform prior. (i) Estimates for homologous boosting parameter, *µ*. (ii) Estimates for homologous boost at the end of the initial waning period, *µ(1 – d)*. (iii) Estimates for duration of initial waning phase, *t*_*s*_. (iv) Estimates for proportion of initial boost lost during the initial waning phase, *d*. (v) Estimates for long term waning rate, *m*. Estimates for TIV 1, TIV 1 + adjuvant and Infection 2 excluded due to lack of identifiability. (vi) Estimates for cross reactivity gradient, *σ*. Note that this value is fixed at 1 for priming infection (Infection 1), shown by the horizontal dotted line. Values for TIV 2 and TIV 1 + adjuvant excluded due to lack of identifiability.

The addition of an adjuvant appeared to have no significant impact on the homologous antibody response to the first vaccine dose, but did improve the response to a second dose of vaccine (TIV 1 compared to TIV 1 + adjuvant and TIV 2 compared TIV 2 + adjuvant, Fig 5ii). Titres against A/Brisbane/10/2007 (H3N2) and A/Solomon Islands/3/2006 (H1N1) were similar following the first unadjuvanted vaccine dose and the first adjuvanted vaccine dose (TIV 1 compared to TIV 1 + adjuvant, Fig 4A&B). However, the second adjuvanted TIV dose appeared to elicit a significant persistent boost to the vaccine strains, which resulted in peak titres near the limit of detection of this assay (TIV 2 compared to TIV 2 + adjuvant, Fig 4A&B).

### Comparison of cross reactivity by exposure type

In models with type-specific cross-reactivity, we found differences in the width of cross reactivity elicited by the 6 exposure types shown in Fig 5. Secondary infection appeared to elicit a level of cross reactivity in line with that of the priming infection, whereas cross reactivity for both unadjuvanted and adjuvanted vaccination appeared to be narrower and only boosted antibodies that were effective against antigenically similar viruses (Fig 6). *σ* describes the degree by which antibody titre decreases as a function of antigenic distance, where higher values of *σ* suggest lower cross reactive breadth. When a single cross reactivity gradient was assumed for all exposure types (as in the highest ranked model), we estimated the cross reactivity gradient to be 2.33 (median; 95% CI:1.74 − 3.01), suggesting narrower cross reactivity than would be expected given the definition for cross reactivity based on ferret antisera (an antigenic distance of 1 unit should see a reduction in antibody boosting of 1 log titre unit). [63] Fig 6 demonstrates that homologous boosting (the y-intercept) was too small to elicit any measurable cross reactive boosting at these antigenic distances. The cross reactivity gradient parameter, *σ*, could therefore not be identified for the second dose of unadjuvanted TIV and first dose of adjuvanted TIV, and we were only able to recover the prior distribution for these parameters. These values were therefore excluded from Fig 5vi.

**Fig 6.**
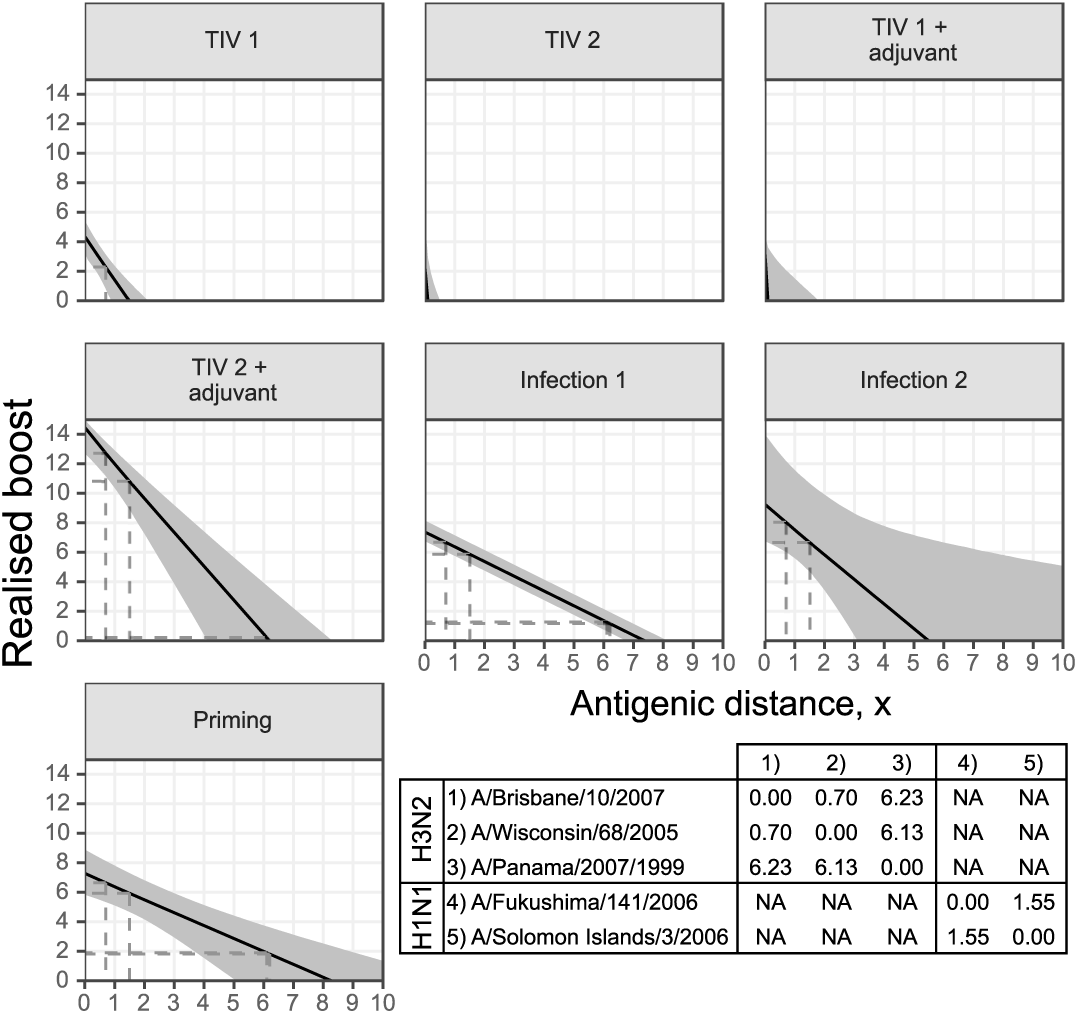
Estimated cross reactivity profiles by exposure type. Solid black lines shows posterior means; shaded regions show 95% credible intervals. Dashed black lines show antigenic distances and corresponding cross reactive boosts given the strains used here. Note that the y-intercept shows the degree of homologous boosting for that exposure type. Table shows assumed antigenic distances between each strain, with no cross reactivity between subtypes.

### Magnitude and duration of waning phases

Our model provided support for the presence of an initial short-term, rapid waning phase followed by a secondary long-term, sustained waning phase. For all vaccine doses, we estimated that the majority of the antibody boost waned within two weeks of reaching the peak (upper 95% CI 17.3, 5.15, 12.5 and 2.16 days for TIV 1, TIV 2, TIV 1 + adjuvant and TIV 2 + adjuvant respectively, Fig 5i & ii). Conversely for priming infection, we estimated that the antibody titre was maintained at near peak levels with an estimated initial waning phase duration of 18.9 days (median; 95% CI 11.0 − 29.3) and a 21.6% (median; 95% CI: 2.02 − 48.8%) drop in log titre relative to the peak. We estimated similar long-term waning rates for second unadjuvanted TIV, second adjuvanted TIV and priming infection (Fig 5v).

### Impact of priming

Prior to receiving non-adjuvanted TIV, experimental group C was infected with H3N2 Panama/2007/1999 at day 0, which represented a host being primed by natural infection prior to vaccination. Our model allowed us to identify additional homologous and cross-reactive antibody boosting that resulted from priming improving the subsequent vaccine response, as comparable experimental groups were given the same vaccination schedule with or without priming infection. We found that priming infection resulted in a substantial additional boost (7.28 log units (median, 95% CI 5.85-8.92)) to antibodies against the A/H1N1 and A/H3N2 vaccine strains at the time of vaccination in addition to that provided by the vaccine itself (Fig 5i).

We estimated the cross reactivity of this additional boost to be broad with a gradient of 0.882 (median; 95% CI: 0.531 − 1.49), suggesting that priming increases the cross-reactive breadth of the vaccine response. It should be noted that whilst additional priming-induced vaccine boosting is well supported by the model fit, the model overestimates the antibody titre to A/Fukushima/141/2006 (H1N1) at day 37 elicited by initial dose of TIV following priming by H3N2 infection (Fig 4C). This may be a result of subtype-specific interactions that are not captured by our model.

### Limited evidence for antigenic seniority and titre dependent boosting

Despite the relatively short duration of these experiments, we found some evidence for a trend of decreasing antibody response with increasing number of prior exposures and/or higher pre-exposure titres. In the best fitting model with antigenic seniority, we estimated *τ* to be 0.213 (median; 95% CI: 0.134-0.300), suggesting that antibody boosting decreased substantially with increasing number exposures after taking into account exposure type and priming. *τ* measures the proportion of the full boost that is lost with each successive exposure experiences relative to the first (ie. boosting decreases linearly as a function of increasing prior exposures). A higher value of *τ* therefore indicates more boosting suppression with an increasing number of prior exposures. Based on these estimates, the amount of antibody boosting would be reduced by over 50% following 4 exposures.

In the best fitting model with titre-dependent boosting, we estimated the titre-dependence gradient *γ* to be 0.0898 (median; 95% CI: 0.0788-0.102) applying to all titres below 10.9 (median; 95% CI: 8.62-11.95). *γ* gives the proportion of full boost that is lost per unit increase in log titre at the time of exposure (with no suppression from a starting log titre of 0). The full posterior estimate for the titre-dependent boosting mechanism is shown in S3 Fig. The top two model variants incorporated one of each of these two mechanisms, suggesting that either one significantly improves model fit relative to the model variants with neither. The two mechanisms are correlated in these experiments, and antigenic seniority was not well identified with estimates for *τ* that did not exclude 0 for models with both titre-dependent boosting and antigenic seniority. However, all of the top models with antigenic seniority but no titre-dependent boosting give constrained estimates for *τ* away from 0 (Fig S12).

## Discussion

In this study, we used a mathematical model of antibody kinetics to describe boosting and waning following influenza vaccination or infection in a group of well characterised ferrets. We fit various subsets of the model with different immunological mechanisms and found that the two best supported models both included: type-specific antibody boosting; type-specific biphasic waning; 6 distinct exposure types; and a role for priming in increasing subsequent vaccine response. Antigenic seniority, antigenic distance-mediated cross reactivity specific to each exposure type and titre-dependent boosting were also included amongst these top models, suggesting that these mechanisms may be important in accurately describing observed antibody titres following multiple exposures. We found quantitative differences in the level of homologous and cross-reactive antibody boosting between vaccination, infection and adjuvanted vaccination in this ferret model. A single TIV dose with or without adjuvant elicited negligible levels of homologous and cross reactive boosting. A second dose of TIV with adjuvant resulted in significant, broadly reactive antibody boosting, whereas a second dose of TIV without adjuvant did not elicit significant antibody boosting. The profile of boosting for primary infection was consistent across experimental groups, and similar in magnitude to secondary infection. Furthermore, we found that priming infection induced a significantly broader and stronger boosting profile following subsequent vaccination.

Our work has a number of limitations. We had insufficient power to quantify all of the mechanisms and parameters proposed here, and therefore restricted our reported results to estimates that were consistent across the best supported model variants (S1 Supporting Protocol). The frequency at which blood samples were taken compared to the number of exposure events resulted in a relatively small amount of data given the number of mechanisms being explored. In particular, sampling around the biphasic waning period of the vaccinations and following the final exposure event was limited, resulting in poor identifiability for some of the waning and timing parameters. Experiments of a similar design with fewer exposures and more frequent sampling would power the model to elucidate these waning phases further and look for differences in response longevity by exposure type.

Our data included only trivalent vaccination with and without Freund’s incomplete adjuvant, and estimates of any boosting parameters are therefore conditional on the presence of three antigens in a single vaccination. It would be interesting to compare the inferred homologous and cross-reactive boost of a three antigen TIV to that elicited by a monovalent vaccine, and for adjuvants more relevant to human vaccination such as MF59. There may also be underlying heterogeneities in antibody response between and within influenza subtypes as well as between vaccine types. [64,65] For example, Live Attenuated Influenza Vaccines (LAIV), as well as newer DNA vaccines may provide different antibody kinetic profiles and may elicit broader antibody responses, or provide different priming effects. [66,67]

We found evidence for biphasic waning following both primary infection and secondary vaccination. There was some evidence that the magnitude and duration of waning differed between exposure types: TIV 1 and 2 and adjuvanted TIV 1 waned very quickly, whereas Infection 2 and TIV 2 + adjuvant were more persistent. Heterogeneity in antibody waning rates between individuals and vaccine types have been shown for other pathogens. [56,68] Although studies of influenza antibody response duration have been carried out in humans, quantifying waning rates independent of subsequent exposures that cause repeated boosting is difficult. [2,69–71] Our model fit to the single exposure ferrets provides an estimation of the waning rate of homologous antibodies in the absence of further exposure, but the cut off of 70 days limits the applicability of this waning rate to a timescale more relevant to humans. Extrapolating our estimated waning rate following primary infection would suggest that antibody titres would wane to non-detectable within a few months, whereas antibody responses against many viruses are known to persist for decades. [68,72] Longer term studies investigating the longevity of the antibody response in the absence of repeated exposure would be useful to quantify a long-term, steady state antibody waning rate. [50] Further mechanisms such as differential waning rates between cross-reactive and homologous antibodies are likely to be important, but were not identifiable here. [21,73] Although animal models are potentially useful, identification of these mechanisms in human populations is likely possible given long-term, frequent sampling of human sera combined with robust statistical methods. [18,22]

Our results have implications for comparing different vaccination strategies. Achieving high HI titres against currently circulating strains is a key endpoint in influenza vaccine trials due to its correlation with clinical protection. [4,74–76] However, there are a number of obstacles to achieving these high titres in some populations including antigenic interactions, age specific responses and antibody waning. [11,77–80] A practical approach to improving vaccine effectiveness is therefore to elicit a broader antibody response to compensate for potential strain mismatch. [81] Adding adjuvants such as MF59 and AS03 has been shown to induce higher antibody titres that have greater cross-reactive properties. [54,55,82,83] Quantitative comparisons of cross reactivity profiles, as we have provided here, could be a useful tool in comparing the effectiveness of different adjuvants, which would provide a measurable benefit to trade-off against safety and immunogenicity concerns. [84,85]

In addition to modelling boosting suppression due to immune imprinting, we considered potential enhancement via priming infection. “Prime-boosting” has been described previously as a strategy to induce broadly reactive immune responses that may be rapidly boosted in advance of exposure to an antigenically novel virus. [51,52,86] Models that included a priming mechanism were ranked systematically higher in our model comparison analysis than those that did not, suggesting that this phenomena is important in explaining titres arising from repeated exposures. We found that vaccine responses to A/H1N1 strains were higher and more broadly reactive in A/H3N2 primed ferrets compared to unprimed ferrets, though our model did not account for subtype specific interactions and subsequently overestimated post vaccination A/H1N1 titres in primed ferrets. Although the phylogenetic relationship between the priming and subsequent boosting strain is likely to be important, heterosubtypic protection has been shown previously in animal models, potentially via cytotoxic T lymphocyte responses. [43,45]

Despite limitations in applying results from animal models directly to humans, these mechanisms may also be relevant to the human immune response by accurately capturing human antibody dynamics at a short time scale. [22] Our results suggest that mathematical models of antibody kinetics that explicitly consider immunological mechanisms and exposure-type specific parameters would be valuable for the prediction of antibody landscapes in human populations. Direct inference from long-term observational data in humans may be difficult, but experimental models, such as the ferret system described here, provide an excellent alternative data source for the inference of short-term immunological mechanisms that may map onto models recovered using human sera. [18,21,39,40]

## Supporting information

S1 Supporting Protocol.Details of the full model, model comparison analysis, model fitting and additional sensitivity analyses.

**S1 Table.Description of model parameters.** Summary of parameter definitions and bounds. All bounds relate to lower and upper bounds of the uniform prior distribution used during model fitting.

S2 Table.Description of model mechanisms and their potential formats.

**S3 Table.Description of models with *δ*ELPD <20.** Table is ranked by ELPD score, such that the model best supported by ELPD (lowest) is at the top.

S4 Table.Summary of parameter estimates for the two best supported models (lowest ELPD score).

S5 Table.csv file containing all posterior distribution estimates for all model variants.

S6 Table.csv file containing convergence diagnostics (including minimum effective sample size and 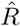) and expected log predictive density estimates for all model variants.

**S1 Fig. Summary of model mechanisms.** A: Cross reactive antibody boosting. The degree of boosting decreases as the antigenic distance between the exposure and measured strain increases. Different exposure types may have different gradients; B: Illustrative example of exposure type specific parameter values. Level of homologous boosting may depend on the exposure type. Note that this may also apply to other parameters eg. waning rate; C: Joint effect of exposure boosting and priming infection. Full boosting following a primed exposure is the sum of contributions of the exposure itself and the effect of priming; D: Antigenic seniority mechanism. Amount of antibody boosting decreases linearly with the number of prior exposures; E: Titre dependent boosting. Solid black line shows example where 0 ≤ *γ* ≤ 1. Blue dashed lines show boundary conditions. Note that the realised boost does not change when *y*_*i*_ is above *y*_*switch*_.

**S2 Fig. Observation error matrix.** Probability of observing a particular log titre given an underlying true, latent titre. Note that the true titre is a continuous value, whereas observations are discrete. Furthermore, truncation of the distribution at the upper and lower limit of the assay results in an asymmetrical distribution when the true value is at either of these limits. True values outside of these limits will be observed as a value within the assay limits.

**S3 Fig. Posterior estimates for titre dependent boosting relationship from the best supported model which included titre dependent boosting.** Shaded gray regions shows 95% credible intervals (CI) drawn from the multivariate posterior. Solid black line shows multivariate posterior mean; Dashed gray lines show median and 95% CI for realised antibody boosting from a titre of 12.

**S4 Fig.Re-estimated model parameters from simulated data.** Violin plots show estimated posterior densities with medians and 95% credible intervals marked as horizontal black lines. Dashed gray lines show bounds on uniform prior. Black dots show true values. (i) Estimates for homologous boosting parameter, *µ*. (ii) Estimates for homologous boost at the end of the initial waning period, *µ(1 – d)*. (iii) Estimates for duration of initial waning phase, *t*_*s*_. (iv) Estimates for proportion of initial boost lost during the initial waning phase, *d*. (v) Estimates for long term waning rate, *m*. Estimates for TIV 1, TIV 1 + adjuvant and Infection 2 excluded due to lack of identifiability. (vi) Estimates for cross reactivity gradient, *σ*. Note that this value is fixed at 1 for priming infection (Infection 1), shown by the horizontal dotted line. Values for TIV 2 and TIV 1 + adjuvant excluded due to lack of identifiability.

**S5 Fig. Summary of posterior distribution estimates for homologous boosting parameter, *µ* from models with *δ*ELPD < 20.** Points show posterior median; line ranges show 95% credible intervals. Estimates are stratified by exposure type and ordered in order of increasing ELPD. Estimates are coloured according to whether or not cross reactivity was assumed to be a universal parameter or type-specific. Dashed horizontal lines represent uniform prior range. Model codes on x-axis relate to the first letter of each mechanism as described in the table.

**S6 Fig. Summary of posterior distribution estimates for initial waning phase proportion, *d* from models with *δ*ELPD < 20.** Points show posterior median; line ranges show 95% credible intervals. Estimates are stratified by exposure type and ordered in order of increasing ELPD. Estimates are coloured according to whether or not titre-dependent boosting was included. Dashed horizontal lines represent uniform prior range. Model codes on x-axis relate to the first letter of each mechanism as described in the table.

**S7 Fig. Summary of posterior distribution estimates for duration of initial waning phase, *t*_*s*_ from models with *δ*ELPD < 20.** Points show posterior median; line ranges show 95% credible intervals. Estimates are stratified by exposure type and ordered in order of increasing ELPD. Estimates are coloured according to whether or not titre-dependent boosting was included. Dashed horizontal lines represent uniform prior range. Model codes on x-axis relate to the first letter of each mechanism as described in the table.

**S8 Fig. Summary of posterior distribution estimates for long-term waning rate, *m* from models with *δ*ELPD < 20.** Points show posterior median; line ranges show 95% credible intervals. Estimates are stratified by exposure type and ordered in order of increasing ELPD. Estimates are coloured according to whether or not waning was assumed to be biphasic or monophasic. Dashed horizontal lines represent uniform prior range. Model codes on x-axis relate to the first letter of each mechanism as described in the table.

**S9 Fig. Summary of posterior distribution estimates for cross reactivity gradient, *σ* from models with *δ*ELPD < 20.** Points show posterior median; line ranges show 95% credible intervals. Estimates are stratified by exposure type and ordered in order of increasing WAIC. Estimates are coloured according to whether or not cross reactivity was assumed to be a universal parameter or type-specific. Plots are truncated from above at 10 for clarity, but upper prior bound was 100. Red dashed line shows the fixed value of *σ* = 1 for priming infection. Blue dashed line shows value above which a homologous boost of µ=5 would give an observed boost of 0 against a strain with an antigenic distance of 1. Model codes on x-axis relate to the first letter of each mechanism as described in the table.

**S10 Fig. Summary of posterior distribution estimates for priming cross reactivity gradient, *β* from models with *δ*ELPD < 20.** Points show posterior median; line ranges show 95% credible intervals. Red dashed line shows the fixed value of *σ* = 1 for priming infection. Blue dashed line shows value above which a homologous boost of µ=5 would give an observed boost of 0 against a strain with an antigenic distance of 1. Estimates are ordered by increasing ELPD. Model codes on x-axis relate to the first letter of each mechanism as described in the table.

**S11 Fig. Summary of posterior distribution estimates for titre dependence gradient, *γ* and titre dependent switch point, *y*_*switch*_ from models with *δ*ELPD < 20.** Points show posterior median; line ranges show 95% credible intervals. Estimates are ordered by increasing ELPD. Model codes on x-axis relate to the first letter of each mechanism as described in the table.

**S12 Fig. Summary of posterior distribution estimates for antigenic seniority parameter, *τ* from models with *δ*ELPD < 20.** Points show posterior median; line ranges show 95% credible intervals. Estimates are ordered by increasing ELPD. Estimates are coloured according to whether or not titre-dependent boosting was also included in the model. Model codes on x-axis relate to the first letter of each mechanism as described in the table.

## Supporting information

S1 Supporting Protocol

## References

1. Kreijtz JHCM, Fouchier RAM, Rimmelzwaan GF. Immune responses to influenza virus infection. Virus Research. 2011;162(1-2):19–30. doi:10.1016/J.VIRUSRES.2011.09.022.

2. Miller E, Hoschler K, Hardelid P, Stanford E, Andrews N, Zambon M. Incidence of 2009 pandemic influenza A H1N1 infection in England: a cross-sectional serological study. Lancet. 2010;375(9720):1100–1108. doi:10.1016/S0140-6736(09)62126-7.

3. Sridhar S, Begom S, Bermingham A, Hoschler K, Adamson W, Carman W, et al. Cellular immune correlates of protection against symptomatic pandemic influenza. Nature Medicine. 2013;19(10):1305–1312. doi:10.1038/nm.3350.

4. Hobson D, Curry RL, Beare AS, Ward-Gardner A. The role of serum haemagglutination-inhibiting antibody in protection against challenge infection with influenza A2 and B viruses. J Hyg. 1972;70(04):767.

5. Potter CW, Oxford JS. Determinants of immunity to influenza infection in man. British medical bulletin. 1979;35(1):69–75.

6. Black S, Nicolay U, Vesikari T, Knuf M, Del Giudice G, Della Cioppa G, et al. Hemagglutination Inhibition Antibody Titers as a Correlate of Protection for Inactivated Influenza Vaccines in Children. The Pediatric Infectious Disease Journal. 2011;30(12):1081–1085. doi:10.1097/INF.0b013e3182367662.

7. Yuan HY, Baguelin M, Kwok KO, Arinaminpathy N, van Leeuwen E, Riley S. The impact of stratified immunity on the transmission dynamics of influenza. Epidemics. 2017;20:84–93. doi:10.1016/j.epidem.2017.03.003.

8. Bedford T, Riley S, Barr IG, Broor S, Chadha M, Cox NJ, et al. Global circulation patterns of seasonal influenza viruses vary with antigenic drift. Nature. 2015;523(7559):217–220. doi:10.1038/nature14460.

9. Russell CA, Jones TC, Barr IG, Cox NJ, Garten RJ, Gregory V, et al. The Global Circulation of Seasonal Influenza A (H3N2) Viruses. Science. 2008;320(5874).

10. Petrova VN, Russell CA. The evolution of seasonal influenza viruses. Nature Reviews Microbiology. 2017;16(1):47–60. doi:10.1038/nrmicro.2017.118.

11. Smith DJ, Forrest S, Ackley DH, Perelson AS. Variable efficacy of repeated annual influenza vaccination. Proc Natl Acad Sci U S A. 1999;96(24):14001–14006.

12. Flannery B, Chung JR, Belongia EA, McLean HQ, Gaglani M, Murthy K, et al. Interim Estimates of 2017–18 Seasonal Influenza Vaccine Effectiveness — United States, February 2018. MMWR Morbidity and Mortality Weekly Report. 2018;67(6):180–185. doi:10.15585/mmwr.mm6706a2.

13. Zost SJ, Parkhouse K, Gumina ME, Kim K, Diaz Perez S, Wilson PC, et al. Contemporary H3N2 influenza viruses have a glycosylation site that alters binding of antibodies elicited by egg-adapted vaccine strains. Proc Natl Acad Sci U S A. 2017;114(47):12578–12583. doi:10.1073/pnas.1712377114.

14. Thompson CP, Lourenço J, Walters AA, Obolski U, Edmans M, Palmer DS, et al. A naturally protective epitope of limited variability as an influenza vaccine target. Nature Communications. 2018;9(1):3859. doi:10.1038/s41467-018-06228-8.

15. Huber VC, Thomas PG, McCullers JA. A multi-valent vaccine approach that elicits broad immunity within an influenza subtype. Vaccine. 2009;27(8):1192–200. doi:10.1016/j.vaccine.2008.12.023.

16. Li GM, Chiu C, Wrammert J, McCausland M, Andrews SF, Zheng NY, et al. Pandemic H1N1 influenza vaccine induces a recall response in humans that favors broadly cross-reactive memory B cells. Proceedings of the National Academy of Sciences of the United States of America. 2012;109(23):9047–52. doi:10.1073/pnas.1118979109.

17. Pebody R, Warburton F, Ellis J, Andrews N, Potts A, Cottrell S, et al. Effectiveness of seasonal influenza vaccine for adults and children in preventing laboratory-confirmed influenza in primary care in the United Kingdom: 2015/16 end-of-season results. Eurosurveillance. 2016;21(38):30348. doi:10.2807/1560-7917.ES.2016.21.38.30348.

18. Kucharski AJ, Lessler J, Read JM, Zhu H, Jiang CQ, Guan Y, et al. Estimating the Life Course of Influenza A(H3N2) Antibody Responses from Cross-Sectional Data. PLoS Biology. 2015;13(3):1–16. doi:10.1371/journal.pbio.1002082.

19. Andrews SF, Huang Y, Kaur K, Popova LI, Ho IY, Pauli NT, et al. Immune history profoundly affects broadly protective B cell responses to influenza. Sci Transl Med. 2015;7(316):316ra192–316ra192. doi:10.1126/scitranslmed.aad0522.

20. Quinñones-Parra SM, Clemens EB, Wang Z, Croom HA, Kedzierski L, McVernon J, et al. A Role of Influenza Virus Exposure History in Determining Pandemic Susceptibility and CD8+ T Cell Responses. Journal of Virology. 2016;90(15):6936–6947. doi:10.1128/JVI.00349-16.

21. Fonville JM, Wilks SH, James SL, Fox A, Ventresca M, Aban M, et al. Antibody landscapes after influenza virus infection or vaccination. Science. 2014;346(6212):7–9.

22. Kucharski AJ, Lessler J, Cummings DAT, Riley S. Timescales of influenza A/H3N2 antibody dynamics. PLoS Biology. 2018;16(8):e2004974. doi:10.1371/journal.pbio.2004974.

23. Wrammert J, Smith K, Miller J, Langley WA, Kokko K, Larsen C, et al. Rapid cloning of high-affinity human monoclonal antibodies against influenza virus. Nature. 2008;453(7195):667–671. doi:10.1038/nature06890.

24. Cobey S, Hensley SE. Immune history and influenza virus susceptibility. Current Opinion in Virology. 2017;doi:10.1016/j.coviro.2016.12.004.

25. Skowronski DM, Chambers C, De Serres G, Sabaiduc S, Winter AL, Dickinson JA, et al. Serial Vaccination and the Antigenic Distance Hypothesis: Effects on Influenza Vaccine Effectiveness During A(H3N2) Epidemics in Canada, 2010-2011 to 2014-2015. J Infect Dis. 2017;215(7):1059–1099.

26. Guthmiller JJ, Wilson PC. Harnessing immune history to combat influenza viruses. Current Opinion in Immunology. 2018;53:187–195. doi:10.1016/J.COI.2018.05.010.

27. Strindhall J, Ernerudh J, Mörner A, Waalen K, Löfgren S, Matussek A, et al. Humoral response to influenza vaccination in relation to pre-vaccination antibody titres, vaccination history, cytomegalovirus serostatus and CD4/CD8 ratio. Infectious Diseases. 2016;48(6):436–442. doi:10.3109/23744235.2015.1135252.

28. Woolthuis RG, Wallinga J, van Boven M. Variation in loss of immunity shapes influenza epidemics and the impact of vaccination. BMC Infectious Diseases. 2017;17(1):632. doi:10.1186/s12879-017-2716-y.

29. Kim JH, Skountzou I, Compans R, Jacob J. Original Antigenic Sin Responses to Influenza Viruses. The Journal of Immunology. 2009;183(5):3294–3301. doi:10.4049/jimmunol.0900398.

30. Gostic KM, Ambrose M, Worobey M, Lloyd-Smith JO. Potent protection against H5N1 and H7N9 influenza via childhood hemagglutinin imprinting. Science. 2016;354(6313).

31. Lessler J, Riley S, Read JM, Wang S, Zhu H, Smith GJD, et al. Evidence for antigenic seniority in influenza A (H3N2) antibody responses in southern China. PLoS Pathogens. 2012;8(7):e1002802. doi:10.1371/journal.ppat.1002802.

32. Lee HY, Topham DJ, Park SY, Hollenbaugh J, Treanor J, Mosmann TR, et al. Simulation and prediction of the adaptive immune response to influenza A virus infection. Journal of virology. 2009;83(14):7151–65. doi:10.1128/JVI.00098-09.

33. Sasaki S, He XS, Holmes TH, Dekker CL, Kemble GW, Arvin AM, et al. Influence of Prior Influenza Vaccination on Antibody and B-Cell Responses. PLoS ONE. 2008;3(8):e2975. doi:10.1371/journal.pone.0002975.

34. Miao H, Hollenbaugh JA, Zand MS, Holden-Wiltse J, Mosmann TR, Perelson AS, et al. Quantifying the early immune response and adaptive immune response kinetics in mice infected with influenza A virus. Journal of virology. 2010;84(13):6687–98. doi:10.1128/JVI.00266-10.

35. Horns F, Vollmers C, Dekker CL, Quake SR. Signatures of selection in the human antibody repertoire: Selective sweeps, competing subclones, and neutral drift. Proc Natl Acad Sci U S A. 2019; p. 201814213. doi:10.1073/pnas.1814213116.

36. Huang KYA, Li CKF, Clutterbuck E, Chui C, Wilkinson T, Gilbert A, et al. Virus-Specific Antibody Secreting Cell, Memory B-cell, and Sero-Antibody Responses in the Human Influenza Challenge Model. The Journal of Infectious Diseases. 2014;209(9):1354–1361. doi:10.1093/infdis/jit650.

37. Angeletti D, Gibbs JS, Angel M, Kosik I, Hickman HD, Frank GM, et al. Defining B cell immunodominance to viruses. Nature Immunology. 2017;18(4):456–463. doi:10.1038/ni.3680.

38. Ferdman J, Palladino G, Liao HX, Moody AM, Kepler TB, Del Giudice G, et al. Intra-seasonal antibody repertoire analysis of a subject immunized with an MF59®-adjuvanted pandemic 2009 H1N1 vaccine. Vaccine. 2018;36(35):5325–5332. doi:10.1016/j.vaccine.2018.06.054.

39. Ranjeva S, Subramanian R, Fang VJ, Leung GM, Ip DKM, Perera RAPM, et al. Age-specific differences in the dynamics of protective immunity to influenza. bioRxiv. 2018; p. 330720. doi:10.1101/330720.

40. Zhao X, Ning Y, Chen MIC, Cook AR. Individual and Population Trajectories of Influenza Antibody Titers Over Multiple Seasons in a Tropical Country. American Journal of Epidemiology. 2018;187(1):135–143. doi:10.1093/aje/kwx201.

41. Laurie KL, Carolan La, Middleton D, Lowther S, Kelso A, Barr IG. Multiple infections with seasonal influenza A virus induce cross-protective immunity against A(H1N1) pandemic influenza virus in a ferret model. The Journal of infectious diseases. 2010;202(7):1011–1020. doi:10.1086/656188.

42. Maher JA, Ms JD. The Ferret : An Animal Model to Study Influenza Virus. Lab animal. 2004;33(9):50–53. doi:10.1038/laban1004-50.

43. McLaren C, Potter CW. Immunity to influenza in ferrets. VII. Effect of previous infection with heterotypic and heterologous influenza viruses on the response of ferrets to inactivated influenza virus vaccines. J Hyg. 1974;72(1):91–100. doi:10.1017/S0022172400023251.

44. Kim JH, Liepkalns J, Reber AJ, Lu X, Music N, Jacob J, et al. Prior infection with influenza virus but not vaccination leaves a long-term immunological imprint that intensifies the protective efficacy of antigenically drifted vaccine strains. Vaccine. 2016;34(4):495–502. doi:10.1016/j.vaccine.2015.11.077.

45. Kreijtz JHCM, Bodewes R, van Amerongen G, Kuiken T, Fouchier RAM, Osterhaus ADME, et al. Primary influenza A virus infection induces cross-protective immunity against a lethal infection with a heterosubtypic virus strain in mice. Vaccine. 2007;25(4):612–620. doi:10.1016/j.vaccine.2006.08.036.

46. Kosikova M, Li L, Radvak P, Ye Z, Wan XF, Xie H. Imprinting of Repeated Influenza A/H3 Exposures on Antibody Quantity and Antibody Quality: Implications on Seasonal Vaccine Strain Selection and Vaccine Performance. Clinical Infectious Diseases. 2018;doi:10.1093/cid/ciy327.

47. Fonville JM, Fraaij PLA, de Mutsert G, Wilks SH, van Beek R, Fouchier RAM, et al. Antigenic Maps of Influenza A(H3N2) Produced With Human Antisera Obtained After Primary Infection. The Journal of infectious diseases. 2016;213(1):31–8. doi:10.1093/infdis/jiv367.

48. Le D, Miller JD, Ganusov VV. Mathematical modeling provides kinetic details of the human immune response to vaccination. Frontiers in Cellular and Infection Microbiology. 2015;4:177. doi:10.3389/fcimb.2014.00177.

49. Halliley JL, Kyu S, Kobie JJ, Walsh EE, Falsey AR, Randall TD, et al. Peak frequencies of circulating human influenza-specific antibody secreting cells correlate with serum antibody response after immunization. Vaccine. 2010;28(20):3582–3587. doi:10.1016/J.VACCINE.2010.02.088.

50. Andraud M, Lejeune O, Musoro JZ, Ogunjimi B, Beutels P, Hens N. Living on Three Time Scales: The Dynamics of Plasma Cell and Antibody Populations Illustrated for Hepatitis A Virus. PLoS Computational Biology. 2012;8(3):e1002418. doi:10.1371/journal.pcbi.1002418.

51. Goji N, Nolan C, Hill H, Wolff M, Noah D, Williams T, et al. Immune Responses of Healthy Subjects to a Single Dose of Intramuscular Inactivated Influenza A/Vietnam/1203/2004 (H5N1) Vaccine after Priming with an Antigenic Variant. The Journal of Infectious Diseases. 2008;198(5):635–641. doi:10.1086/590916.

52. Rudenko L, Naykhin A, Donina S, Korenkov D, Petukhova G, Isakova-Sivak I, et al. Assessment of immune responses to H5N1 inactivated influenza vaccine among individuals previously primed with H5N2 live attenuated influenza vaccine. Hum Vaccin Immunother. 2015;11(12):2839–2848.

53. Bodewes R, Kreijtz JHCM, Geelhoed-Mieras MM, van Amerongen G, Verburgh RJ, van Trierum SE, et al. Vaccination against seasonal influenza A/H3N2 virus reduces the induction of heterosubtypic immunity against influenza A/H5N1 virus infection in ferrets. Journal of virology. 2011;85(6):2695–702. doi:10.1128/JVI.02371-10.

54. van den Brand JMA, Kreijtz JHCM, Bodewes R, Stittelaar KJ, van Amerongen G, Kuiken T, et al. Efficacy of vaccination with different combinations of MF59-adjuvanted and nonadjuvanted seasonal and pandemic influenza vaccines against pandemic H1N1 (2009) influenza virus infection in ferrets. Journal of virology. 2011;85(6):2851–8. doi:10.1128/JVI.01939-10.

55. Khurana S, Chearwae W, Castellino F, Manischewitz J, King LR, Honorkiewicz A, et al. Vaccines with MF59 adjuvant expand the antibody repertoire to target protective sites of pandemic avian H5N1 influenza virus. Science translational medicine. 2010;2(15):15ra5. doi:10.1126/scitranslmed.3000624.

56. White M, Idoko O, Sow S, Diallo A, Kampmann B, Borrow R, et al. Antibody kinetics following vaccination with MenAfriVac: an analysis of serological data from randomised trials. The Lancet Infectious Diseases. 2019;0(0). doi:10.1016/S1473-3099(18)30674-1.

57. Amanna IJ, Slifka MK. Mechanisms that determine plasma cell lifespan and the duration of humoral immunity. Immunological Reviews. 2010;236(1):125–138. doi:10.1111/j.1600-065X.2010.00912.x.

58. Plummer, M., Best, N., Cowles, K., and Vines, K. coda: Output analysis and diagnostics for MCMC. R News. 2008;6.1:7–11 Available from: https://cran.r-project.org/package=coda.

59. Vehtari A, Gabry J, Yao Y, Gelman A. loo: Efficient leave-one-out cross-validation and WAIC for Bayesian models; 2018. Available from: https://cran.r-project.org/package=loo.

60. Vehtari A, Gelman A, Gabry J. Practical Bayesian model evaluation using leave-one-out cross-validation and WAIC. Statistics and Computing. 2017;27(5):1413–1432. doi:10.1007/s11222-016-9696-4.

61. Yao Y, Vehtari A, Simpson D, Gelman A. Using stacking to average Bayesian predictive distributions. Bayesian Analysis. 2017;doi:10.1214/17-BA1091.

62. Galipaud M, Gillingham MAF, David M, Dechaume-Moncharmont FX. Ecologists overestimate the importance of predictor variables in model averaging: a plea for cautious interpretations. Methods in Ecology and Evolution. 2014;5(10):983–991. doi:10.1111/2041-210X.12251.

63. Smith DJ, Lapedes AS, de Jong JC, Bestebroer TM, Rimmelzwaan GF, Osterhaus ADME, et al. Mapping the antigenic and genetic evolution of influenza virus. Science. 2004;305(5682):371–6. doi:10.1126/science.1097211.

64. Freeman G, Pereram RAPM, Ngan E, Fang VJ, Cauchemez S, Ip DKM, et al. Quantifying homologous and heterologous antibody titre rises after influenza virus infection. Epidemiology and Infection. 2016;144(11):2306–2316. doi:10.1017/S0950268816000583.

65. Knudsen NPH, Olsen A, Buonsanti C, Follmann F, Zhang Y, Coler RN, et al. Different human vaccine adjuvants promote distinct antigen-independent immunological signatures tailored to different pathogens. Scientific reports. 2016;6:19570. doi:10.1038/srep19570.

66. Hoft DF, Lottenbach KR, Blazevic A, Turan A, Blevins TP, Pacatte TP, et al. Comparisons of the Humoral and Cellular Immune Responses Induced by Live Attenuated Influenza Vaccine and Inactivated Influenza Vaccine in Adults. Clin Vaccine Immunol. 2017;24(1).

67. Basha S, Hazenfeld S, Brady RC, Subbramanian RA. Comparison of antibody and T-cell responses elicited by licensed inactivated-and live-attenuated influenza vaccines against H3N2 hemagglutinin. Human Immunology. 2011;72(6):463–469. doi:10.1016/j.humimm.2011.03.001.

68. Antia A, Ahmed H, Handel A, Carlson NE, Amanna IJ, Antia R, et al. Heterogeneity and longevity of antibody memory to viruses and vaccines. PLoS Biology. 2018;16(8):e2006601. doi:10.1371/journal.pbio.2006601.

69. Petrie JG, Ohmit SE, Johnson E, Truscon R, Monto AS. Persistence of Antibodies to Influenza Hemagglutinin and Neuraminidase Following One or Two Years of Influenza Vaccination. Journal of Infectious Diseases. 2015;212(12):1914–1922. doi:10.1093/infdis/jiv313.

70. Wright PF, Sannella E, Shi JR, Zhu Y, Ikizler MR, Edwards KM. Antibody Responses After Inactivated Influenza Vaccine in Young Children. The Pediatric Infectious Disease Journal. 2008;27(11):1004–1008. doi:10.1097/INF.0b013e31817d53c5.

71. Frank AL, Taber LH, Glezen WP, Paredes A, Couch RB. Reinfection with Influenza A (H3N2) Virus in Young Children and Their Families. Journal of Infectious Diseases. 1979;140(6):829–833. doi:10.1093/infdis/140.6.829.

72. Amanna IJ, Carlson NE, Slifka MK. Duration of humoral immunity to common viral and vaccine antigens. The New England journal of medicine. 2007;357(19):1903–1915. doi:10.1542/peds.2008-2139LLLL.

73. Jacobson RM, Grill DE, Oberg AL, Tosh PK, Ovsyannikova IG, Poland GA. Profiles of influenza A/H1N1 vaccine response using hemagglutination-inhibition titers. Human Vaccines & Immunotherapeutics. 2015;11(4):961–969. doi:10.1080/21645515.2015.1011990.

74. Reber A, Adrian R, Jacqueline K, Katz J. Immunological assessment of influenza vaccines and immune correlates of protection. Expert review of vaccines. 2013;12(5):519–36. doi:10.1586/erv.13.35.

75. Dormitzer PR, Galli G, Castellino F, Golding H, Khurana S, Del Giudice G, et al. Influenza vaccine immunology. Immunological Reviews. 2011;239(1):167–177. doi:10.1111/j.1600-065X.2010.00974.x.

76. Coudeville L, Bailleux F, Riche B, Megas F, Andre P, Ecochard R. Relationship between haemagglutination-inhibiting antibody titres and clinical protection against influenza: development and application of a bayesian random-effects model. BMC medical research methodology. 2010;10:18. doi:10.1186/1471-2288-10-18.

77. Belongia EA, Sundaram ME, McClure DL, Meece JK, Ferdinands J, VanWormer JJ. Waning vaccine protection against influenza A (H3N2) illness in children and older adults during a single season. Vaccine. 2015;33(1):246–251. doi:10.1016/j.vaccine.2014.06.052.

78. Ferdinands JM, Fry AM, Reynolds S, Petrie J, Flannery B, Jackson ML, et al. Intraseason waning of influenza vaccine protection: Evidence from the US Influenza Vaccine Effectiveness Network, 2011-12 through 2014-15. Clin Infect Dis. 2017;64(5):544–550.

79. Mugitani A, Ito K, Irie S, Eto T, Ishibashi M, Ohfuji S, et al. Immunogenicity of the Trivalent Inactivated Influenza Vaccine in Young Children Less than 4 Years of Age, with a Focus on Age and Baseline Antibodies. Clinical and Vaccine Immunology. 2014;21(9):1253–1260. doi:10.1128/CVI.00200-14.

80. Mosterín Höpping A, McElhaney J, Fonville JM, Powers DC, Beyer WEP, Smith DJ, et al. The confounded effects of age and exposure history in response to influenza vaccination. Vaccine. 2016;34(4):540–546. doi:10.1016/j.vaccine.2015.11.058.

81. Allen JD, Owino SO, Carter DM, Crevar CJ, Reese VA, Fox CB, et al. Broadened immunity and protective responses with emulsion-adjuvanted H5 COBRA-VLP vaccines. Vaccine. 2017;35(38):5209–5216. doi:10.1016/J.VACCINE.2017.07.107.

82. Forrest HL, Khalenkov AM, Govorkova EA, Kim JK, Del Giudice G, Webster RG. Single-and multiple-clade influenza A H5N1 vaccines induce cross protection in ferrets. Vaccine. 2009;27(31):4187–4195. doi:10.1016/j.vaccine.2009.04.050.

83. Jackson LA, Campbell JD, Frey SE, Edwards KM, Keitel WA, Kotloff KL, et al. Effect of Varying Doses of a Monovalent H7N9 Influenza Vaccine With and Without AS03 and MF59 Adjuvants on Immune Response: A Randomized Clinical Trial. JAMA. 2015;314(3):237–246.

84. Shoenfeld Y, Agmon-Levin N. ’ASIA’ – Autoimmune/inflammatory syndrome induced by adjuvants. J Autoimmun. 2011;36(1):4–8.

85. Gupta RK, Relyveld EH, Lindblad EB, Bizzini B, Ben-Efraim S, Gupta CK. Adjuvants–a balance between toxicity and adjuvanticity. Vaccine. 1993;11(3):293–306.

86. Galli G, Hancock K, Hoschler K, DeVos J, Praus M, Bardelli M, et al. Fast rise of broadly cross-reactive antibodies after boosting long-lived human memory B cells primed by an MF59 adjuvanted prepandemic vaccine. Proc Natl Acad Sci U S A. 2009;106(19):7962–7967. doi:10.1073/pnas.0903181106.

