## Supplementary material for "Characterising antibody kinetics from multiple influenza infection and vaccination events in ferrets": S1 Supporting Protocol

#### Materials and Methods

##### Base boosting and biphasic waning model

We defined a deterministic model of antibody kinetics that included additive boosting and biphasic waning, as represented by Fig 1 in the main text. Equation 1 describes a simplified version of the model, where antibody levels to a single strain are described over time following a single exposure to influenza antigens. Here, we define the general term for any kind of influenza antigen exposure (eg. inoculation, vaccination) as an *exposure*. We also define *effective antibodies against strain A* or *measured strain A* as the log HI assay titre measurement that was effective at inhibiting hemagglutination by antigens of influenza strain $A$. Similarly, we use the term *exposure strain* to describe the strain of influenza antigens contained in a given exposure (eg. vaccine strain).

Prior to any form of exposure, the initial antibody titre, $y_i^0$, was undetectable (log titre of 0). After an exposure event at a known time, $\xi$, antibody levels to a particular influenza strain were linearly boosted on the log scale by an amount $\mu$ after a given time interval $t_p$. Titres then quickly dropped by a fixed proportion, $d$, as the short-lived component of the antibody response wanes over time $t_s$. Antibody titres then enter a long-term, slower waning phase at rate $m$ until subsequent exposure, when antibody dynamics become dominated by a new set of boosting and waning parameters. Note that Equation 1 is left expanded to distinguish between parameters as intervals of time ($t_p$, $t_s$), the time of exposure as a fixed point in time ($\xi$) and the time variable, $t$.

$$f(\theta_i, t) = \begin{cases} y_i^0 & t \leq \xi_i \\ \frac{\mu_i}{t_{pi}} t - \frac{\mu_i}{t_{pi}} \xi_i + y_i^0 & \xi_i < t \leq \xi_i + t_{pi} \\ -\frac{d_i \mu_i}{t_{si}} t + \frac{d_i \mu_i}{t_{si}} (\xi_i + t_{pi}) + \mu_i + y_i^0 & \xi_i + t_{pi} < t \leq \xi_i + t_{pi} + t_{si} \\ -m_i t + m_i(\xi_i + t_{pi} + t_{si}) + (1 - d_i)\mu_i + y_i^0 & \xi_i + t_{pi} + t_{si} < t \leq \xi_{(i+1)} \end{cases} \quad (1)$$

Where:

$$\theta_i = \{\mu_i, d_i, t_{pi}, t_{si}, m_i, \xi_i\} \quad (2)$$

$y_i^0$ represents the antibody titre at the time of exposure $i$. Prior to the first exposure, this value is set to 0, representing the absence of any influenza antibodies.

#### Interaction of multiple exposures

Each successive influenza exposure provides a dose of antigen that stimulates the adaptive immune response. An observed trajectory of antibodies over time is therefore the culmination of antibody responses generated from each exposure. Although there may be some interaction between overlapping exposures due to competition for T-helper cell recruitment or interference in binding to antigen, [1, 2] we assumed that all of the exposures here were sufficiently far apart to be effectively independent. [3, 4] In order, the times between exposures were 28 days, 14 days and 28 days. In the model, each exposure elicited boosting and biphasic waning phases starting from the time of that exposure. We assumed that antibody kinetics followed one contiguous process, whereby the dynamics at a particular time are governed only by the parameters of the most recent exposure. In this sense, observed antibody titres are a direct reflection of independent antibody secreting cell (ASC) populations that undergo phases of boosting and waning depending on the time of the most recent exposure. [5] Each subsequent exposure supersedes the previous exposure such that:

$$y_A(t) = \left\{ \begin{array}{ll} f(\theta_1, t) & 0 \leq t < \xi_2 \\ f(\theta_2, t) & \xi_2 \leq t < \xi_3 \\ \vdots & \\ f(\theta_n, t) & \xi_n \leq t \end{array} \right\} \quad (3)$$

$y_i^0$ is defined as:

$$y_i^0 = \left\{ \begin{array}{ll} 0 & i = 1 \\ f(\theta_{i-1}, \xi_i) & i > 1 \end{array} \right\} \quad (4)$$

#### Additional model mechanisms

The most basic form of the multiple exposure model assumes that titres for each measured strain following each exposure have their own unique set of parameters, where $\theta = \{\theta_1, \theta_2, \ldots, \theta_n\}$ for an exposure history with $n$ exposures and $\theta_1 \cap \theta_2 \cap \cdots \cap \theta_n = \emptyset$. Whilst this simple descriptive model may provide good within-sample predictive accuracy, its dependence on a particular set of experiments makes it difficult to generalise, as parameters would need to be estimated for each exposure strain, measured strain, exposure time and exposure type. We therefore sought to add additional structure to the model to describe biological mechanisms that may play a role in shaping an individual's antibody profile. These mechanisms include cross-reactive boosting, exposure-type specific parameters, a role for priming in increased antibody boosting, antigenic seniority and titre-dependent boosting. These model mechanisms are depicted in Fig S1, and their implementation is described below.

#### Biphasic and monophasic waning

Measurement of serum antibody titres after exposure are typically described as a biphasic waning process following peak titre, as short-lived ASCs (that account for the initial boost in titre) begin to die and are succeeded by more persistent antibody-secreting plasma cells. [6] We also considered a simpler model with only a single waning phase by eliminating the initial waning phase. This is achieved by fixing

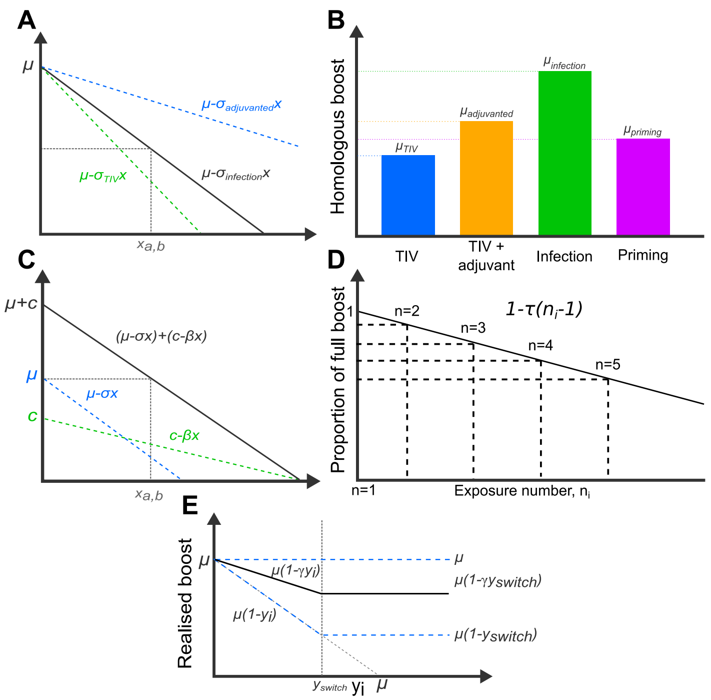


**Fig S1. Summary of model mechanisms** A: Cross reactive antibody boosting. The degree of boosting decreases as the antigenic distance between the exposure and measured strain increases. Different exposure types may have different gradients; B: Illustrative example of exposure type specific parameter values. Level of homologous boosting may depend on the exposure type. Note that this may also apply to other parameters eg. waning rate; C: Joint effect of exposure boosting and priming infection. Full boosting following a primed exposure is the sum of contributions of the exposure itself and the effect of priming; D: Antigenic seniority mechanism. Amount of antibody boosting decreases linearly with the number of prior exposures; E: Titre dependent boosting. Solid black line shows example where $0 \leq \gamma \leq 1$. Blue dashed lines show boundary conditions. Note that the realized boost does not change when $y_i$ is above $y_{switch}$.

$t_s = d = 0.$

#### Cross reactive boosting

A useful property of influenza antibodies is that antibodies produced in response to exposure with a given strain, A, may be partially effective at hemagglutination inhibition against a different strain, B. The strength of this cross reactivity is a function of how antigenically similar A and B are in shape space. [7,8] This property of influenza viruses enables useful phylogenetic and antigenic comparison, such as the inference of antigenic maps or antibody landscapes. [9,10] Given that infection of a naive ferret with strain A elicits a maximum antibody titre to strain A of $\mu$ log HI units, strain B has an antigenic distance from strain A of 1 if the antibody titre to strain B is $\mu - 1$ using sera from the same ferret. We used this property to set the parameter vector $\theta$ to be conditional on the antigenic distance between the exposure strain and the measured strain (Fig S1A). The degree of boosting of antibodies effective against strain A following exposure with strain B, $\mu_{A,B}$, can therefore be generalised as:

$$\mu_{A,B} = \mu' - \sigma x_{A,B} \tag{5}$$

Where $\mu'$ is the level of homologous boosting following any exposure; $\sigma$ is the cross reactivity gradient by which antibody effectiveness drops with increasing antigenic distance; and $x_{A,B}$ is the antigenic distance between strain A and strain B.

The cross reactivity gradient, $\sigma$, for initial infection was fixed at 1 in line with previous definitions of antigenic distance. [7] Estimating the cross reactivity gradient for other exposure types is therefore taken relative to this, such that a gradient of less than 1 suggests broader cross reactivity than priming infection, and greater than 1 suggests narrower cross reactivity. Antigenic distances were assumed to be fixed and known based on euclidean antigenic distance taken from previous data, and no cross reactivity in the HI assay was allowed between subtypes. [11] The antigenic distance between each pair based on previous data was: 6.23 between A/Panama/2007/1999 and A/Brisbane/10/2007; 6.12 between A/Panama/2007/1999 and A/Wisconsin/67/2005; 0.70 between A/Brisbane/10/2007 and A/Wisconsin/67/2005; and 1.55 between A/Fukushima/141/2006 and A/Solomon Islands/3/2006. [10]

#### Exposure types and type-specific parameters

Different influenza exposure types have been shown to generate different antibody boosting and persistence. [12–16] We therefore inferred exposure-specific boosting and waning parameters for each of the exposure types in the protocol. We considered either 3 distinct types (infection, vaccination and adjuvanted vaccination) or 6 distinct exposure type (one for each unique exposure formulation: primary infection; initial TIV; secondary TIV; initial adjuvanted TIV; secondary adjuvanted TIV; secondary infection).

Exposure type, $l$, was defined as:

$$l \in \{\text{infection}, \text{TIV}, \text{TIV} + \text{adjuvant}\} \tag{6}$$

or

$$\begin{aligned} l \in \quad & \{\text{infection}_1,\ \text{infection}_2,\ \text{TIV}_1,\ \text{TIV}_2, \\ & \text{TIV}_1 + \text{adjuvant},\ \text{TIV}_2 + \text{adjuvant}\} \end{aligned} \tag{7}$$

Note that as the number of exposure events in the experiments was far greater than the number of different exposure types, this grouping into exposures types substantially reduces the number of parameters in the model (Fig S1B). We also considered two scenarios for type specific cross reactivity, whereby cross reactivity was either conditional on exposure type, $l$, (ie. $\sigma_l$), or universal across all exposure types (ie. one value of $\sigma$ for all exposures).

#### The effect of priming

Priming by infection has been shown to elicit both an increased magnitude and breadth of antibody boosting following subsequent vaccination. [17, 18] To compare the impact of priming infection on vaccine response, we modified the boosting parameter $\mu$ to consider additional antibody boosting following vaccination if the ferret had previously received a priming infection. We assumed that vaccination following priming infection elicited an additional degree of homologous boosting as well as cross-reactive boosting with a different cross-reactivity gradient compared to un-primed vaccination. The term for antibody boosting including priming was defined as:

$$\mu_{A,B} = (\mu' - \sigma x_{A,B}) + \alpha(c - \beta x_{A,B}) \tag{8}$$

Where $c$ is the magnitude of additional homologous boosting due to priming; $\beta$ is the gradient of cross reactivity related to priming infection (similar to $\sigma$); and $\alpha$ is either 1 or 0 depending on if priming had or had not occurred (Fig S1C).

#### Antigenic seniority

The number of previous exposures has been shown to impact antibody responses following exposure. [19, 20] Evidence for antigenic seniority suggests that the antibody response to each new exposure is smaller than the response from the previous exposure. We included this mechanism in our model by modifying the boosting parameter, $\mu$, to be conditional on the number of previous exposures (Fig S1D):

$$\mu_{A,B,n} = \big(1 - \tau(n-1)\big)\big((\mu' - \sigma x_{A,B}) + \alpha(c - \beta x_{A,B})\big) \tag{9}$$

Where $\tau$ gives the proportion of the full boost lost relative to the first exposure $(0 \leq \tau \leq 1)$ as a function of the number of previous exposures, $n-1$. In these experiments, ferrets may have experienced up to 4 exposures.

#### Titre dependent boosting

Ceiling effects have been observed in influenza antibody responses previously, where individuals already at a high antibody titre exhibit lower levels of boosting compared to naive individuals. [21, 22] We captured this mechanism by modifying the boosting parameter such that:

$$\mu_{A,B,n,y_i(\xi)} = \left\{ \begin{array}{ll} \mu_{A,B,n}(1 - \gamma y_i(\xi)) & y_i(\xi) < y_{switch} \\ \mu_{A,B,n}(1 - \gamma y_{switch}) & y_i(\xi) \geq y_{switch} \end{array} \right\} \tag{10}$$

Where $y_i(\xi)$ indicates the antibody titre against strain $i$ at the time of exposure, $\xi$; $\gamma$ is the degree by which antibody boosting drops with preexisting antibody titre; and $y_{switch}$ is a boundary on $y_i(\xi)$ above which antibody boosting remains unchanged (Fig S1E). $\gamma$ was bounded between 0 and 1, where 0 implies no titre dependence and 1 implies strong titre dependent suppression of boosting.

#### Full model

Incorporating all of the mechanisms above gives the full set of model parameters for a given exposure:

$$\theta_{i,A,B,l} = \{\mu'_l, d_l, t_{pl}, t_{sl}, m_l, \xi_i, x_{A,B}, \sigma_l, \alpha_i, c, \beta, \gamma, y_{switch}, \tau\} \tag{11}$$

The realized level of antibody boosting to a particular strain is given by:

$$\mu_{i,A,B,n,l,y_i(\xi)} = \big(1 - \gamma y_i(\xi)\big)\big(1 - \tau(n-1)\big)\big((\mu' - \sigma x_{A,B}) + \alpha(c - \beta x_{A,B})\big) \tag{12}$$

Where $i$ is the exposure index (specifying only exposure time and whether the exposure was primed); $A$ is the measured strain; $B$ is the exposure strain; $n-1$ is the number of previous exposures; and $l$ is the type of exposure. All model parameters are described in Table S1. Note that the indicated subscript for each parameter is crucial in highlighting where parameters are shared between different exposure events.

It is possible to include or exclude all of these mechanisms by fixing or removing certain parameters: setting $t_s = d = 0$ incorporates monophasic waning, biphasic waning otherwise; cross reactivity may be either universal or type specific; typing may include either 3 or 6 distinct exposure types; fixing $\tau$ to 0 removes antigenic seniority; titre dependent boosting can be removed by setting $\gamma = 0$; and fixing $c$ to 0 removes priming. The result is 6 mechanisms each with 2 settings, giving 64 different model combinations to be fitted (Table S2).

#### Model fitting

The model predicts log antibody titres as latent states on a continuous scale for a given set of parameter values. We assumed that observed antibody titres followed a truncated normal distribution with mean given by the model predicted latent titre. HI titres were taken as the highest 2-fold dilution of sera at which hemagglutination was inhibited, and the observed value was therefore discrete and observed as the lower integer bound (ie. a true log titre of 5.5 would be observed as 5). Furthermore, the limits of the assay meant that only log values between 0 and 12 could be observed. The likelihood of

**Table S1.** Description of model parameters. Summary of parameter definitions and bounds. All bounds relate to lower and upper bounds of the uniform prior distribution used during model fitting.

| Parameter Name | Definition | Lower bound | Upper bound | Notes |
|---|---|---|---|---|
| $\mu$ | Homologous boost on log scale | 0 | 15 | |
| $t_p$ | Time from exposure to peak titre (days) | 12 | 12 | Fixed at 12 days |
| $d$ | Proportion of boost waned in initial waning | 0 | 1 | Fixed at 0 for monophasic waning |
| $t_s$ | Duration of initial waning phase | 0 | 30 | Fixed at 0 for monophasic waning |
| $m$ | Long term titre waning rate (log units per day) | 0 | 12 | |
| $\sigma$ | Cross reactivity gradient | 0 | 100 | A value above 5 would give a cross reactive boost of <0.1 log units at an antigenic distance of 0.5 |
| $\tau$ | Antigenic seniority modifier | 0 | 1 | Antigenic seniority modified given by $\big(1-\tau(n-1)\big)$ where $n-1$ is the number of previous exposures |
| $\gamma$ | Titre dependence gradient | 0 | 1 | 0 gives no titre dependence. 1 gives strong titre dependent suppression of antibody boosting from higher titres |
| $y_{switch}$ | Maximum titre at which titre dependent boosting is in effect | 0 | 12 | Initial titres above this have the same titre dependence as y_switch |
| $\beta$ | Priming cross reactivity gradient | 0 | 100 | A value above 5 would give a cross reactive boost of <0.1 log units at an antigenic distance of 0.5 |
| $c$ | Additional boost due to priming | 0 | 15 | Fixed at 0 to remove priming |
| $\epsilon$ | Standard deviation of truncated normal distribution | 0 | 10 | |

observing a particular titre, $k$, given a true titre, $y$, is given by:

$$L(k|y,\epsilon) = \begin{cases} f(x<1) & \text{if } k=0 \\ f(k \leq x < k+1) & \text{if } 1 \leq k < 12 \\ f(x \geq 12) & \text{if } k \geq 12 \end{cases} \tag{13}$$

Where $\epsilon$ is the standard deviation of the normal distribution; and $f$ is the cumulative distribution function of the standard normal distribution given by:

$$f(x) = \Phi(\frac{x-y}{\epsilon}) = \frac{1}{2}\left[1+\operatorname{erf}\left(\frac{x-y}{\epsilon\sqrt{2}}\right)\right] \tag{14}$$

**Table S2.** Description of model mechanisms and their potential formats.

| Mechanism | Option 1 | Option 2 |
|---|---|---|
| Waning | Biphasic | Monophasic |
| Cross reactivity | Type specific | Universal |
| Typed exposures | 3 types | 6 types |
| Priming | Present | Absent |
| Antigenic seniority | Present | Absent |
| Titre dependent boosting | Present | Absent |

Where $erf(x)$ is the error function, given by:

$$\mathrm{erf}(x) = \frac{2}{\sqrt{\pi}} \int_0^x e^{-t^2} dt \tag{15}$$

Using this probability function, we defined an observation error matrix giving the probability of an observed log HI titre given a true, underlying log titre, as shown in Fig S2.

We sampled from the multivariate posterior distribution of the parameter vector, $\theta$, which is comprised of different parameters depending on the included model mechanisms. We assumed uniform prior distributions for all free model parameters, with upper and lower bounds for these priors described in the Table S1. Parameter estimates were obtained for each set of $\mu$, $d$, $t_s$ and $m$, as well as $\sigma$, $\beta$, $c$, $\rho$, $\gamma$ and $y_{switch}$. The time to peak titre parameter, $t_p$, was fixed at 12 days. We took 5000000 samples from the multivariate posterior using parallel-tempering Markov chain Monte Carlo (PT-MCMC) to achieve an effective sample size (ESS) of at least 200 for all parameters. Convergence was assessed visually and based on Gelman-Rubin convergence diagnostics with the 'coda' R package. Where the ESS was $< 200$ or the upper 95% confidence interval on the potential scale reduction factor was $> 1.1$ for any parameter, chains were re-run for 10000000 and with a greater number of temperatures to improve convergence. 1 of the 1296 estimated parameters still had an upper 95% confidence interval for $\hat{R} = 1.103$ and another had an ESS of 197, though neither of these were presented in main text estimates (only 5 other parameters had an ESS $< 400$). All analyses were performed with R and C++. All code and data are available as an R package at https://github.com/jameshay218/antibodyKinetics.

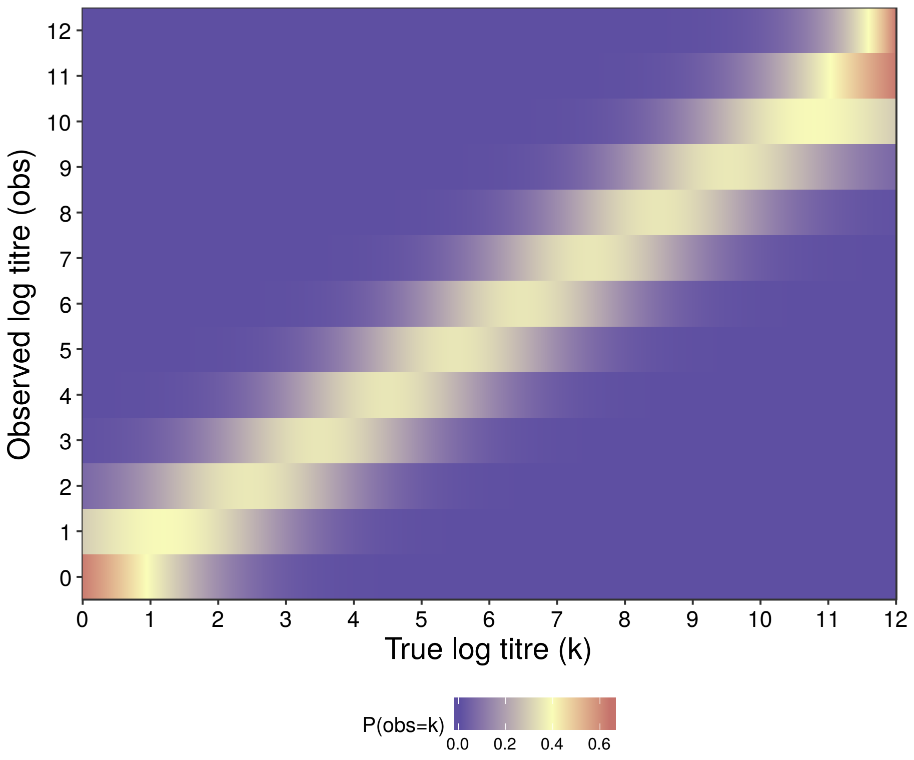


**Fig S2.** Observation error matrix. Probability of observing a particular log titre given an underlying true, latent titre. Note that the true titre is a continuous value, whereas observations are discrete. Furthermore, truncation of the distribution at the upper and lower limit of the assay results in an asymmetrical distribution when the true value is at either of these limits. True values outside of these limits will be observed as a value within the assay limits.

#### Model fitting and comparison

To validate the base model before incorporating additional complexity arising from multiple exposures, we first fit the model for a single exposure to a single virus from group E, where 3 ferrets were only infected with a single H3N2 virus (A/Panama/2007/99 (H3N2)). We compared four variants of this base model to justify the inclusion of full biphasic waning: boosting followed by no antibody waning ($t_s = d = m = 0$); boosting followed by monophasic waning ($t_s = d = 0$); boosting followed by biphasic waning with no longer term waning ($m = 0$); and boosting followed by biphasic waning (all model parameters free). We then fit various iterations of the full model to the complete data set (5 groups each of 3 ferrets) incorporating each combination of mechanisms, ranging from the most biologically simple model with 8 free parameters (monophasic waning; type non-specific cross reactivity; no antigenic seniority, priming effect or titre-dependent boosting; and 3 distinct exposure types) to the most complex with 35 free parameters (biphasic waning; type-specific cross-reactivity; antigenic seniority; titre-dependent boosting; priming; and 6 distinct exposure types). The time to peak titre parameter, $t_p$, was fixed at 12 days for all exposures due to the lack of data immediately after each exposure. We also performed a sensitivity analysis varying $t_p$ parameter.

We then added each mechanism described above into the model such that we could compare model fits for each additional level of biological complexity. Note that biological complexity here (more mechanisms) is not the same as model complexity (number of free parameters), as more biological mechanisms add additional structure across experimental groups to restrict parameter space at the cost of model flexibility. By ranking models on their information criteria (which takes into account model fit penalised by model complexity), we aimed to (i) justify the inclusion of more complex mechanisms in our model and (ii) check for consistency of parameter estimates between model variants. We compared models based on their expected log pointwise predictive density (ELPD), estimated using Pareto-smoothed importance sampling leave-one-out cross-validation (PSIS-LOO) with the 'loo' R package, and the widely applicable information criterion (WAIC). PSIS-LOO and WAIC serve a similar purpose to the Akiake Information Criterion (AIC) but are more suitable in a Bayesian setting. [23–25] Mechanisms that were more common amongst higher ranked models are better supported by the data, whereas mechanisms that are absent do not provide sufficient additional explanatory power to justify their added complexity to the model. Our aim was not to maximize predictive performance but rather to quantify mechanisms of immunological interest and justify interpretation of estimated parameter values. We therefore did not perform a full Bayesian model averaging analysis or formal variable selection, though we did perform Pseudo-Bayesian model averaging (Pseudo-BMA+) to estimate the relative weights of each model variant and therefore each mechanism.

When using PSIS-LOO, the 'loo' package calculates the shape parameter, $\hat{k}$, for each data point for each fitted model. For many of the model variants, $\hat{k}$ estimates were $> 0.7$ for a small number of data points ( 1%), suggesting that the PSIS-LOO estimated ELPD estimates were not reliable for these fits. Visual inspection suggested that problematic data points were outliers that were not adequately described by the models or were highly influential. We refit each model excluding each data point with $\hat{k} > 0.7$ (leave one out cross-validation) to directly calculate the ELPD for these data points, and updated our model ELPD estimates accordingly.

Finally, we performed simulation recovery experiments for the base model and all model variants with a $\delta$ELPD$<$20 to test that we could accurately estimate known model parameters.

### Additional Results

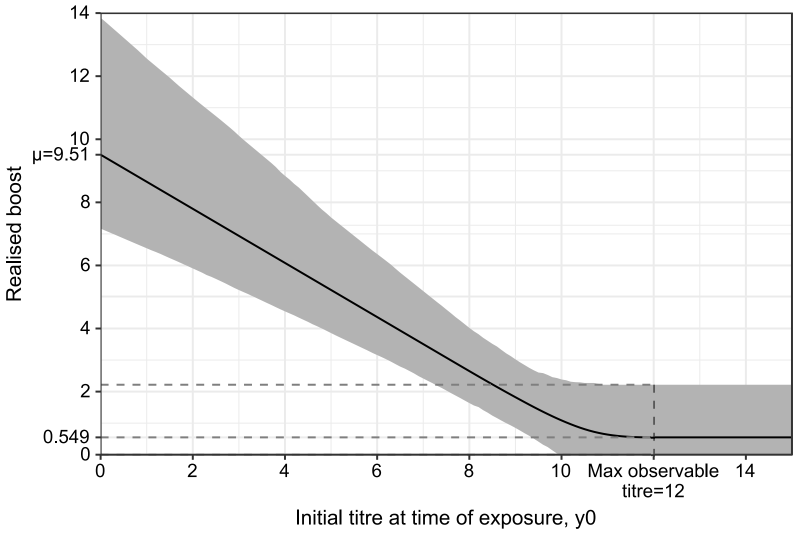


**Fig S3. Posterior estimates for titre dependent boosting relationship from the best supported model which included titre dependent boosting.** Shaded gray regions shows 95% credible intervals (CI) drawn from the multivariate posterior. Solid black line shows multivariate posterior mean; Dashed gray lines show median and 95% CI for realised antibody boosting from a titre of 12.

#### Simulation studies

We performed a simulation-recovery experiment to test that our model fitting framework was able to re-estimate known model parameters from simulated data that matched the experimental protocol for group E (3 individuals, infection on day 0, HI titres measured at days 0, 21, 37, 49 and 70). 95% credible intervals (CI) for the marginal posteriors of the model parameters all encompassed the true parameter values, though were wide for some parameters. These were: true value $\mu = 10.0$, estimated $\mu = 8.49$ (median, 95%CI 6.36-12.4); true value $d = 0.5$, estimated $m = 0.499$ (median, 95%CI 0.128-0.666); true value $t_s = 19$ days, estimated $t_s = 18.0$ days (median, 95% CI 3.16-27.7 days); true value $m = 0.04$, estimated $m = 0.0609$ (median, 95%CI 0.0207-0.105). Performing simulation-recovery on simulated data with weekly HI titres resulted in more constrained marginal posteriors, suggesting that further experiments with more frequent observations would provide more tightly constrained estimates: true value $\mu = 10.0$, estimated $\mu = 9.47$ (median, 95%CI 8.76-10.2); true value $d = 0.5$, estimated $d = 0.628$ (median, 95%CI 0.506-0.715); true value $t_s = 19$ days, estimated $t_s = 24.5$ days (median,

95% CI 17.2-27.8 days); true value $m = 0.04$, estimated $m = 0.0428$ (median, 95%CI 0.00898-0.0777). There was some bias in the inferred boosting and initial waning proportion towards sharper initial waning. This may be attributed to the discretised nature of the data, where all observations of the hidden, continuous true antibody titre are observed as the lower integer. For example, all true log titres $5 \leq x < 6$ would be observed as a log titre of 5, therefore the posterior probability for any combined value of $5 \leq d\mu < 6$ would be uniform given one observed log HI titre $k = 5$ at time $t_p + t_s$.

We also ran simulation recovery experiments for the 13 models with a $\delta$ELPD$< 20$ to test that we could estimate parameters using simulated data generated from the experimental protocol. We took the maximum likelihood parameter values from the real model fits and used these to simulate data under each of these 13 models that matched data from the real experimental protocol (5 groups of 3 ferrets, 5 tested strains, 5 blood samples taken). We considered a parameter re-estimation as accurate when the 95% and 99% credible intervals (CI) of the inferred posterior distribution encompassed the true parameter value. Under this definition, the parameter reestimation accuracy was 87.5% (median; range 73.3-100%) for the 95% CI case and 90.6% (median; range 83.3%-100%). The vast majority of inaccurately estimated parameters either narrowly missed the true value or were the result of very weak identification that recovered only the prior for that parameter. Fig S4 shows the inferred posterior distributions against the true parameter values for the model variant with biphasic waning, 6 exposure types, priming, antigenic seniority, type specific cross reactivity and no titre-dependent boosting. Although posterior estimates were weakly constrained for some parameters as in the real data, the models fit to and explained the simulated observations well.

###### 0.0.1 Further model comparison results

Performing direct leave-one-out cross validation for the data points with $\hat{k} > 0.7$ resulted in the fitting of a further 375 models. We applied the same convergence diagnostics for these runs as in the main text (5000000 iterations, minimum effective sample size of 200 and $\hat{R} < 1.1$ for all parameters). Where either of these conditions were violated, we re-ran the MCMC chains for 10000000 iterations. After running the chains for 10000000 iterations, the upper 95% confidence interval for $\hat{R}$ was between 1.1 and 1.3 for 26 of the 7891 estimated parameters, suggesting that there was still some

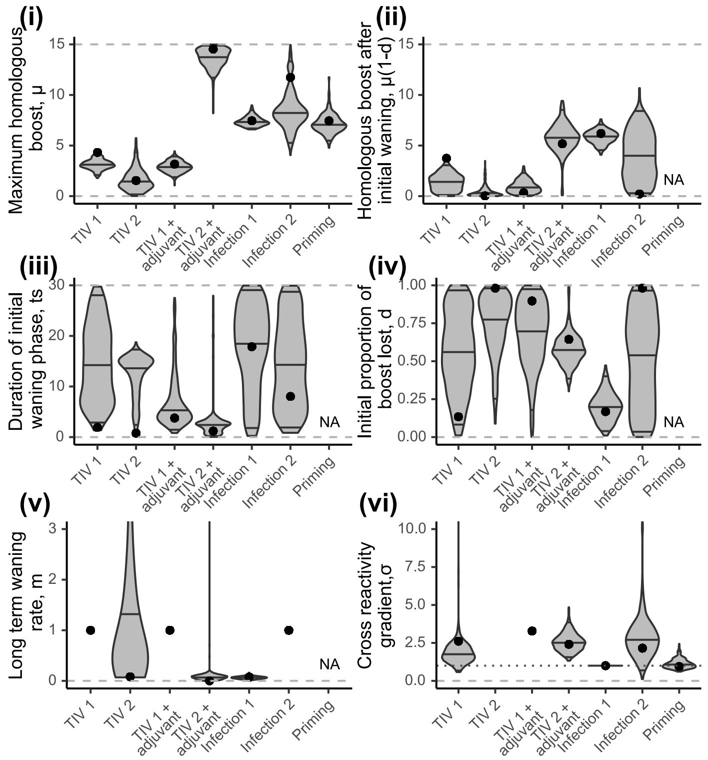


**Fig S4. Re-estimated model parameters from simulated data.** Violin plots show estimated posterior densities with medians and 95% credible intervals marked as horizontal black lines. Dashed gray lines show bounds on uniform prior. Black dots show true values. (i) Estimates for homologous boosting parameter, $\mu$. (ii) Estimates for homologous boost at the end of the initial waning period, $\mu(1 - d)$. (iii) Estimates for duration of initial waning phase, $t_s$. (iv) Estimates for proportion of initial boost lost during the initial waning phase, $d$. (v) Estimates for long term waning rate, $m$. Estimates for TIV 1, TIV 1 + adjuvant and Infection 2 excluded due to lack of identifiability. (vi) Estimates for cross reactivity gradient, $\sigma$. Note that this value is fixed at 1 for priming infection (Infection 1), shown by the horizontal dotted line. Values for TIV 2 and TIV 1 + adjuvant excluded due to lack of identifiability.

between-chain variance in the inferred posterior mean for 0.3% of the parameters. Inspection of each of these marginal posteriors revealed that this lack of convergence was due to poorly identified parameters, where changes in parameter value have no impact on model predicted values. For example, the majority of problematic parameters were long-term waning rates ($m$) where the majority of probability density was at parameter values $m < 1$, but with some remaining density spread uniformly across $1 < m < 12$. Given that we are interested in comparing the model predicted values to the observed data in this step and not in obtaining accurate parameter estimates, we were confident that these 26 parameters would not impact our ELPD estimates.

A linear regression with ELPD as the outcome variable and included mechanisms as predictors suggested that inclusion of antigenic seniority, priming, 6 exposure types, biphasic waning and titre-dependent boosting were all associated with improved model fits, but inclusion type-specific cross reactivity was not. Regression coefficients were: presence of antigenic seniority, 6.51 (95% CI: 1.05-12.0); priming, 77.7 (95%CI: 72.3-83.2); 6 exposure types compared to 3 exposure types, 21.7 (95% CI: 16.3-27.2); biphasic waning compared to monophasic waning, 11.4 (95%CI: 5.97-16.9); titre-dependent boosting, 13.3 (95%CI: 7.82-18.7); no type-specific cross reactivity, 2.26 (95% CI: -3.19-7.73).

#### Sensitivity analyses

##### Consistency of parameter estimates across model variants

Parameter estimates for comparable mechanisms were broadly consistent across the 13 best supported model variants with a $\delta$ELPD$<20$ compared to the best fitting model (Table S3). Parameter estimates for all 64 model variants are given in Table S5. Across the model variants with 6 distinct exposure types and biphasic waning, estimates for boosting and initial waning parameters were similar within each exposure type (Fig S5, Fig S6 and Fig S7. Models with $\delta$ELPD of less than 20 are shown). None of the best supported model variants included only three exposure types (unadjuvanted TIV, adjuvanted TIV and infection), though those that did suggested low levels of transient boosting following TIV and TIV + adjuvant, and high levels of persistent boosting following infection. These results contrast with the consistent estimates of higher levels of persistent boosting from TIV 2 + adjuvant for the rest of the model variants, which were better able to describe the differences in boosting observed between the first and second vaccine doses. Similarly, all of the 13 models included a role for priming infection in increasing subsequent vaccine response. For model variants with identifiable biphasic waning, estimates for the long-term waning rate were consistently low across all exposure types, though generally higher for TIV 1, TIV 1 + adjuvant and Infection 2 (Fig S8). 5 of the models with a $\delta$ELPD$<20$ included monophasic waning, suggesting that monophasic waning was able to accurately describe observed antibody titres from these experiments.

All but 2 of the top 13 models included antigenic seniority and/or titre-dependent boosting, suggesting that models accounting for a pattern of decreasing antibody boosting improved predictive power (Fig S12). Estimates for type specific cross-reactivity parameters were also consistent across model variants; vaccination elicited an antigenically narrower response than priming infection, whereas secondary infection and additional primed-vaccine boosting elicited a cross-reactive response similar to that of priming infection (Fig S9 and Fig S10).

##### Varying time to peak parameter

Varying the time to peak antibody titre parameter, $t_p$, did not significantly change the inferred parameter estimates. We re-fit the top 3 models (based on ELPD) under the two scenarios: a) the time to peak titre was considered to be unknown, but constrained between 10 and 14 days for primary infection and between 5 and 14 days for all other exposures; b) the time to peak titre was fixed at 12 days for primary infection and 8 days for all other exposures. In all cases, the ELPD of each model variant was highest with $t_p$ fixed at 12 days for all exposures, as in the main results. Estimating $t_p$ or fixing $t_p$ at lower values resulted in poor identifiability of parameter $t_s$. However, the inferred parameter estimates for all other parameters were very similar whether $t_p$ was fixed or estimated, suggesting that this assumption did not bias our results.

#### References


1. Zarnitsyna VI, Ellebedy AH, Davis C, Jacob J, Ahmed R, Antia R. Masking of antigenic epitopes by antibodies shapes the humoral immune response to influenza. Philosophical transactions of the Royal Society of London Series B, Biological sciences. 2015;370(1676):20140248–. doi:10.1098/rstb.2014.0248.

2. Zarnitsyna VI, Lavine J, Ellebedy AH, Ahmed R, Antia R. Multi-epitope Models Explain How Pre-existing Antibodies Affect the Generation of Broadly Protective Responses to Influenza. PLOS Pathogens. 2016;12(6):e1005692. doi:10.1371/journal.ppat.1005692.

3. Cao P, Yan AWC, Heffernan JM, Petrie S, Moss RG, Carolan LA, et al. Innate Immunity and the Inter-exposure Interval Determine the Dynamics of Secondary Influenza Virus Infection and Explain Observed Viral Hierarchies. PLoS Computational Biology. 2015;11(8):e1004334. doi:10.1371/journal.pcbi.1004334.

4. Laurie KL, Guarnaccia TA, Carolan LA, Yan AWC, Aban M, Petrie S, et al. Interval Between Infections and Viral Hierarchy Are Determinants of Viral Interference Following Influenza Virus Infection in a Ferret Model. The Journal of infectious diseases. 2015;212(11):1701–10. doi:10.1093/infdis/jiv260.

5. Amanna IJ, Slifka MK. Mechanisms that determine plasma cell lifespan and the duration of humoral immunity. Immunological Reviews. 2010;236(1):125–138. doi:10.1111/j.1600-065X.2010.00912.x.

6. Lam JH, Baumgarth N. The Multifaceted B Cell Response to Influenza Virus. The Journal of Immunology. 2019;202(2):351–359. doi:10.4049/jimmunol.1801208.

7. Smith DJ, Lapedes AS, de Jong JC, Bestebroer TM, Rimmelzwaan GF, Osterhaus ADME, et al. Mapping the antigenic and genetic evolution of influenza virus. Science. 2004;305(5682):371–6. doi:10.1126/science.1097211.

8. Lapedes A, Farber R. The Geometry of Shape Space: Application to Influenza. Journal of Theoretical Biology. 2001;212(1):57–69. doi:10.1006/jtbi.2001.2347.

9. Fonville JM, Wilks SH, James SL, Fox A, Ventresca M, Aban M, et al. Antibody landscapes after influenza virus infection or vaccination. Science. 2014;346(6212):7–9.

10. Bedford T, Suchard MA, Lemey P, Dudas G, Gregory V, Hay AJ, et al. Integrating influenza antigenic dynamics with molecular evolution. eLife. 2014;2014(3):e01914. doi:10.7554/eLife.01914.

11. Burlington DB, Wright PF, van Wyke KL, Phelan MA, Mayner RE, Murphy BR. Development of subtype-specific and heterosubtypic antibodies to the influenza A virus hemagglutinin after primary infection in children. J Clin Microbiol. 1985;21(5):847–849.

12. Wright PF, Sannella E, Shi JR, Zhu Y, Ikizler MR, Edwards KM. Antibody Responses After Inactivated Influenza Vaccine in Young Children. The Pediatric Infectious Disease Journal. 2008;27(11):1004–1008. doi:10.1097/INF.0b013e31817d53c5.

13. Hoft DF, Lottenbach KR, Blazevic A, Turan A, Blevins TP, Pacatte TP, et al. Comparisons of the Humoral and Cellular Immune Responses Induced by Live Attenuated Influenza Vaccine and Inactivated Influenza Vaccine in Adults. Clin Vaccine Immunol. 2017;24(1).

14. Kang EK, Lim JS, Lee JA, Kim DH. Comparison of immune response by virus infection and vaccination to 2009 pandemic influenza A/H1N1 in children. Journal of Korean medical science. 2013;28(2):274–9. doi:10.3346/jkms.2013.28.2.274.

15. Sridhar S, Brokstad K, Cox R. Influenza Vaccination Strategies: Comparing Inactivated and Live Attenuated Influenza Vaccines. Vaccines. 2015;3(2):373–389. doi:10.3390/vaccines3020373.

16. Vella S, Rocchi G, Resta S, Marcelli M, De Felici A. Antibody reactive in antibody-dependent cell-mediated cytotoxicity following influenza virus vaccination. Journal of medical virology. 1980;6(3):203–11.

17. Goji N, Nolan C, Hill H, Wolff M, Noah D, Williams T, et al. Immune Responses of Healthy Subjects to a Single Dose of Intramuscular Inactivated Influenza A/Vietnam/1203/2004 (H5N1) Vaccine after Priming with an Antigenic Variant. The Journal of Infectious Diseases. 2008;198(5):635–641. doi:10.1086/590916.

18. Rudenko L, Naykhin A, Donina S, Korenkov D, Petukhova G, Isakova-Sivak I, et al. Assessment of immune responses to H5N1 inactivated influenza vaccine among individuals previously primed with H5N2 live attenuated influenza vaccine. Hum Vaccin Immunother. 2015;11(12):2839–2848.

19. Lessler J, Riley S, Read JM, Wang S, Zhu H, Smith GJD, et al. Evidence for antigenic seniority in influenza A (H3N2) antibody responses in southern China. PLoS Pathogens. 2012;8(7):e1002802. doi:10.1371/journal.ppat.1002802.

20. Mosterín Höpping A, McElhaney J, Fonville JM, Powers DC, Beyer WEP, Smith DJ, et al. The confounded effects of age and exposure history in response to influenza vaccination. Vaccine. 2016;34(4):540–546. doi:10.1016/j.vaccine.2015.11.058.

21. Jacobson RM, Grill DE, Oberg AL, Tosh PK, Ovsyannikova IG, Poland GA. Profiles of influenza A/H1N1 vaccine response using hemagglutination-inhibition titers. Human Vaccines & Immunotherapeutics. 2015;11(4):961–969. doi:10.1080/21645515.2015.1011990.

22. Petrie JG, Ohmit SE, Johnson E, Cross RT, Monto AS. Efficacy studies of influenza vaccines: effect of end points used and characteristics of vaccine failures. J Infect Dis. 2011;203(9):1309–1315. doi:10.1093/infdis/jir015.

23. Vehtari A, Gabry J, Yao Y, Gelman A. loo: Efficient leave-one-out cross-validation and WAIC for Bayesian models; 2018. Available from: `https://cran.r-project.org/package=loo`.

24. Gelman A, Hwang J, Vehtari A. Understanding predictive information criteria for Bayesian models. Stat Comput. 2013;24(6):997–1016.

25. Vehtari A, Gelman A, Gabry J. Practical Bayesian model evaluation using leave-one-out cross-validation and WAIC. Statistics and Computing. 2017;27(5):1413–1432. doi:10.1007/s11222-016-9696-4.

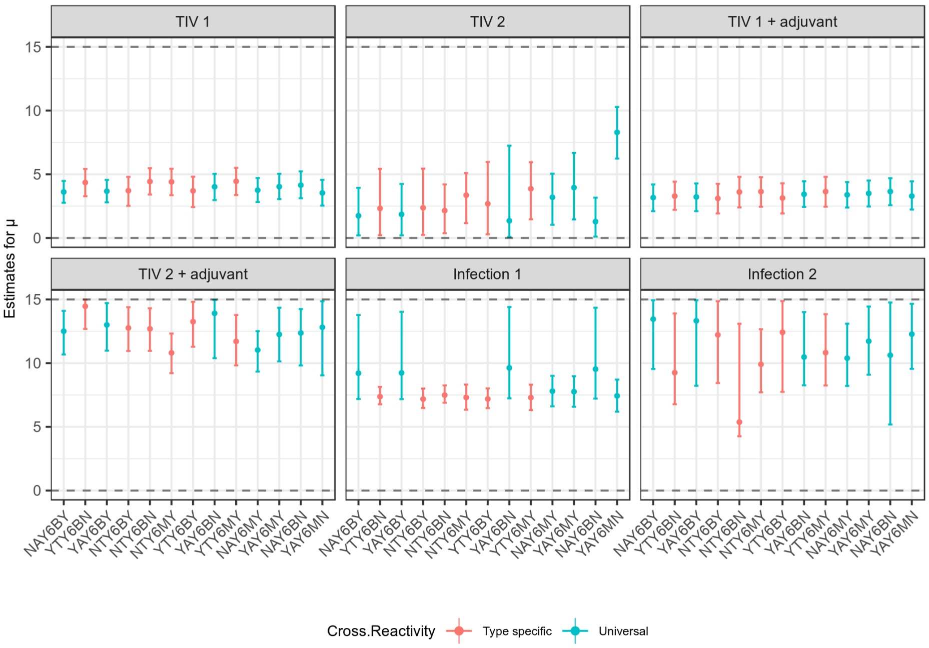


**Fig S5. Summary of posterior distribution estimates for homologous boosting parameter, $\mu$ from models with $\delta$ELPD $< 20$.** Points show posterior median; line ranges show 95% credible intervals. Estimates are stratified by exposure type and ordered in order of increasing ELPD. Estimates are coloured according to whether or not cross reactivity was assumed to be a universal parameter or type-specific. Dashed horizontal lines represent uniform prior range. Model codes on x-axis relate to the first letter of each mechanism as described in the table.

**Table S4. Summary of parameter estimates for the two best supported models (lowest ELPD score).**

**Table S5. .csv file containing all posterior distribution estimates for all model variants.**

**Table S6. .csv file containing convergence diagnostics (including minimum effective sample size and $\hat{R}$) and expected log predictive density estimates for all model variants.**

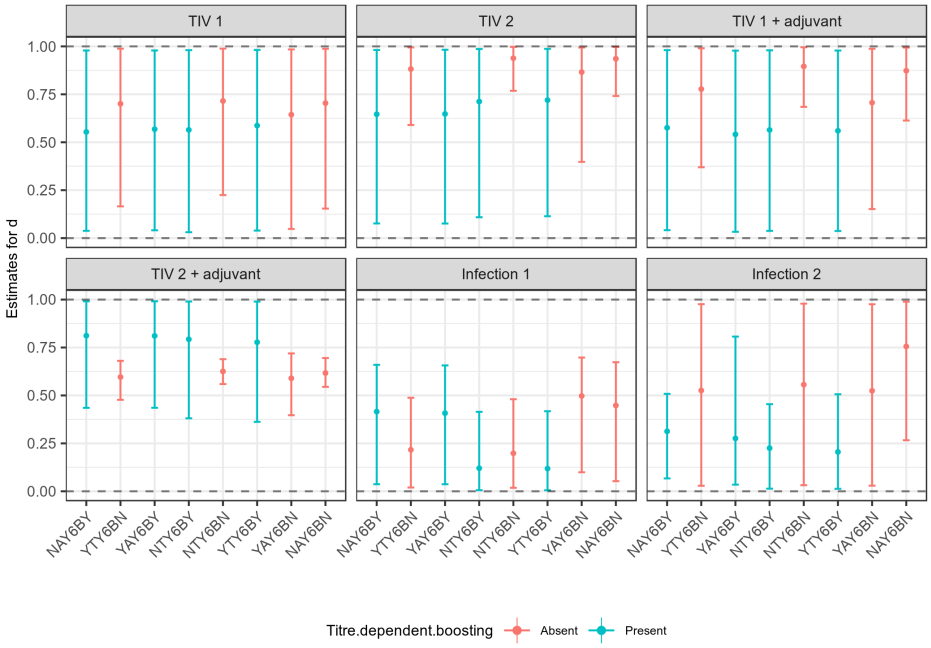


**Fig S6. Summary of posterior distribution estimates for initial waning phase proportion, $d$ from models with $\delta$ELPD $< 20$.** Points show posterior median; line ranges show 95% credible intervals. Estimates are stratified by exposure type and ordered in order of increasing ELPD. Estimates are coloured according to whether or not titre-dependent boosting was included. Dashed horizontal lines represent uniform prior range. Model codes on x-axis relate to the first letter of each mechanism as described in the table.

**Table S3.** Description of models with $\delta$ELPD <20. Table is ranked by ELPD score, such that the model best supported by ELPD (lowest) is at the top.

| **Run Identifier** | **Antigenic Seniority** | **Cross Reactivity** | **Priming** | **Typed exposures** | **Waning** | **Titre dependent boosting** | **$\delta$ELPD** | **ELPD standard error** |
|---|---|---|---|---|---|---|---|---|
| NAY6BY | Absent | Universal | Present | 6 types | Biphasic | Present | 0 | 21.373 |
| YTY6BN | Present | Type specific | Present | 6 types | Biphasic | Absent | 0.421 | 20.656 |
| YAY6BY | Present | Universal | Present | 6 types | Biphasic | Present | 2.059 | 21.279 |
| NTY6BY | Absent | Type specific | Present | 6 types | Biphasic | Present | 2.636 | 21.194 |
| NTY6BN | Absent | Type specific | Present | 6 types | Biphasic | Absent | 4.06 | 20.599 |
| NTY6MY | Absent | Type specific | Present | 6 types | Monophasic | Present | 4.625 | 20.938 |
| YTY6BY | Present | Type specific | Present | 6 types | Biphasic | Present | 4.83 | 21.172 |
| YAY6BN | Present | Universal | Present | 6 types | Biphasic | Absent | 5.185 | 21.146 |
| YTY6MY | Present | Type specific | Present | 6 types | Monophasic | Present | 5.807 | 20.889 |
| NAY6MY | Absent | Universal | Present | 6 types | Monophasic | Present | 7.781 | 21.135 |
| YAY6MY | Present | Universal | Present | 6 types | Monophasic | Present | 7.996 | 21.131 |
| NAY6BN | Absent | Universal | Present | 6 types | Biphasic | Absent | 10.53 | 21.036 |
| YAY6MN | Present | Universal | Present | 6 types | Monophasic | Absent | 17.607 | 20.841 |

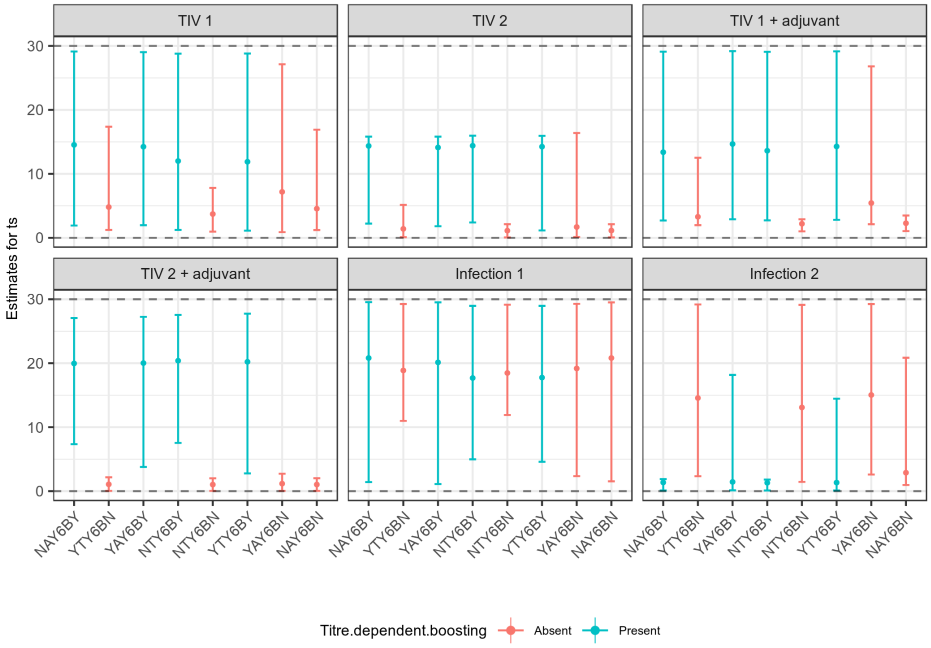


**Fig S7. Summary of posterior distribution estimates for duration of initial waning phase, $t_s$ from models with $\delta$ELPD $< 20$.** Points show posterior median; line ranges show 95% credible intervals. Estimates are stratified by exposure type and ordered in order of increasing ELPD. Estimates are coloured according to whether or not titre-dependent boosting was included. Dashed horizontal lines represent uniform prior range. Model codes on x-axis relate to the first letter of each mechanism as described in the table.

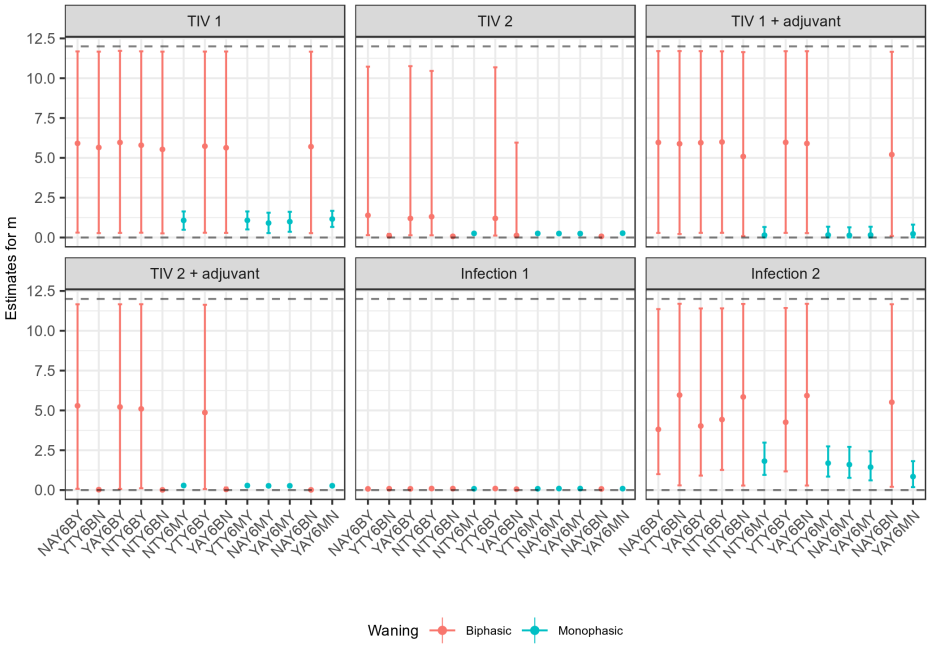


**Fig S8. Summary of posterior distribution estimates for long-term waning rate, $m$ from models with $\delta$ELPD $< 20$.** Points show posterior median; line ranges show 95% credible intervals. Estimates are stratified by exposure type and ordered in order of increasing ELPD. Estimates are coloured according to whether or not waning was assumed to be biphasic or monophasic. Dashed horizontal lines represent uniform prior range. Model codes on x-axis relate to the first letter of each mechanism as described in the table.

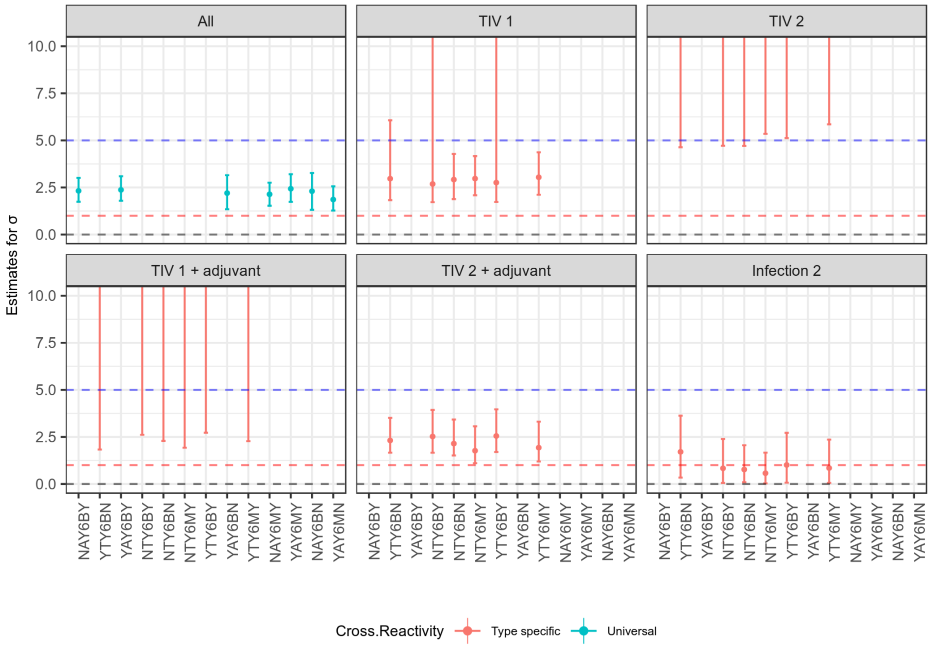


**Fig S9. Summary of posterior distribution estimates for cross reactivity gradient, $\sigma$ from models with $\delta$ELPD $< 20$.** Points show posterior median; line ranges show 95% credible intervals. Estimates are stratified by exposure type and ordered in order of increasing ELPD. Estimates are coloured according to whether or not cross reactivity was assumed to be a universal parameter or type-specific. Plots are truncated from above at 10 for clarity, but upper prior bound was 100. Red dashed line shows the fixed value of $\sigma = 1$ for priming infection. Blue dashed line shows value above which a homologous boost of µ=5 would give an observed boost of 0 against a strain with an antigenic distance of 1. Model codes on x-axis relate to the first letter of each mechanism as described in the table.

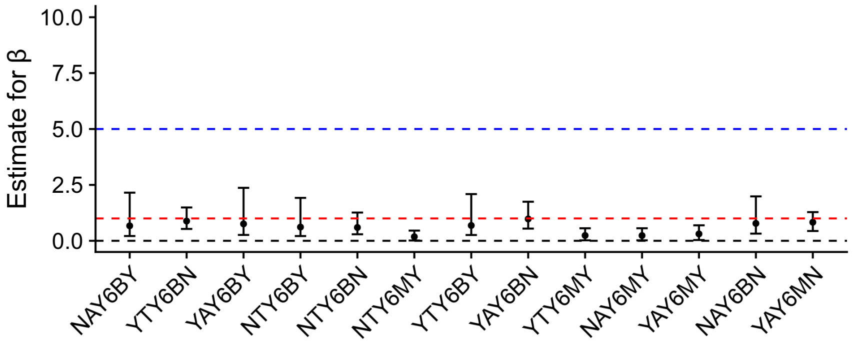


**Fig S10. Summary of posterior distribution estimates for priming cross reactivity gradient, $\beta$ from models with $\delta$ELPD $< 20$.** Points show posterior median; line ranges show 95% credible intervals. Red dashed line shows the fixed value of $\sigma = 1$ for priming infection. Blue dashed line shows value above which a homologous boost of µ=5 would give an observed boost of 0 against a strain with an antigenic distance of 1. Estimates are ordered by increasing ELPD. Model codes on x-axis relate to the first letter of each mechanism as described in the table.

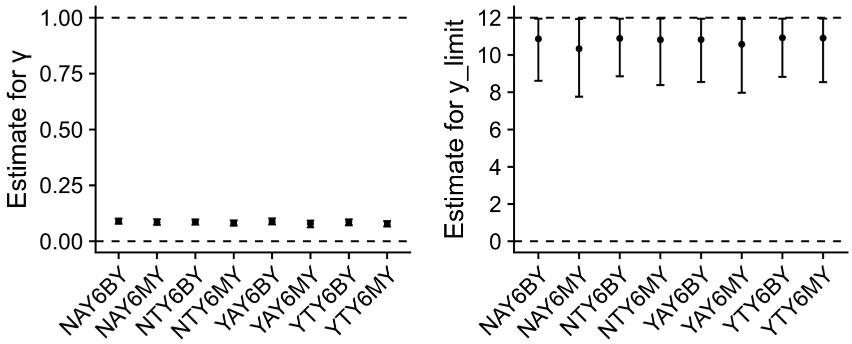


**Fig S11. Summary of posterior distribution estimates for titre dependence gradient, $\gamma$ and titre dependent switch point, $y_{switch}$ from models with $\delta$ELPD $< 20$.** Points show posterior median; line ranges show 95% credible intervals. Estimates are ordered by increasing ELPD. Model codes on x-axis relate to the first letter of each mechanism as described in the table.

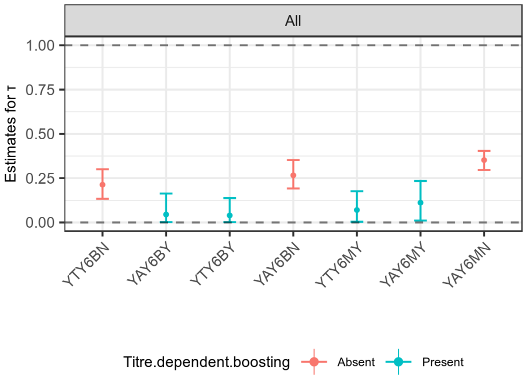


**Fig S12. Summary of posterior distribution estimates for antigenic seniority parameter, $\tau$ from models with $\delta$ELPD $< 20$.** Points show posterior median; line ranges show 95% credible intervals. Estimates are ordered by increasing ELPD. Estimates are coloured according to whether or not titre-dependent boosting was also included in the model. Model codes on x-axis relate to the first letter of each mechanism as described in the table.
